# Genetically encoded fluorescent reporter for polyamines

**DOI:** 10.1101/2024.08.24.609500

**Authors:** Pushkal Sharma, Colin Y. Kim, Heather R. Keys, Shinya Imada, Alex B. Joseph, Luke Ferro, Tenzin Kunchok, Rachel Anderson, Omer Yilmaz, Jing-Ke Weng, Ankur Jain

**Affiliations:** Whitehead Institute of Biomedical Research, Cambridge, MA, USA; Department of Chemical Engineering, Massachusetts Institute of Technology, Cambridge, MA, USA; Department of Biological Engineering, Massachusetts Institute of Technology, Cambridge, MA, USA; Department of Molecular and Cellular Biology, Harvard University, Cambridge, MA, USA; The David H. Koch Institute for Integrative Cancer Research at MIT, Cambridge, MA, USA; Department of Biology, Massachusetts Institute of Technology, Cambridge, MA, USA; Institute for Plant-Human Interface, Northeastern University, Boston, MA, USA; Department of Chemistry and Chemical Biology, Department of Bioengineering and Department of Chemical Engineering, Northeastern University, Boston, MA, USA

## Abstract

Polyamines are abundant and evolutionarily conserved metabolites that are essential for life. Dietary polyamine supplementation extends life-span and health-span. Dysregulation of polyamine homeostasis is linked to Parkinson’s disease and cancer, driving interest in therapeutically targeting this pathway. However, measuring cellular polyamine levels, which vary across cell types and states, remains challenging. We introduce a first-in-class genetically encoded polyamine reporter for real-time measurement of polyamine concentrations in single living cells. This reporter utilizes the polyamine-responsive ribosomal frameshift motif from the OAZ1 gene. We demonstrate broad applicability of this approach and reveal dynamic changes in polyamine levels in response to genetic and pharmacological perturbations. Using this reporter, we conducted a genome-wide CRISPR screen and uncovered an unexpected link between mitochondrial respiration and polyamine import, which are both risk factors for Parkinson’s disease. By offering a new lens to examine polyamine biology, this reporter may advance our understanding of these ubiquitous metabolites and accelerate therapy development.

## Introduction

Polyamines are small aliphatic compounds with two or more amine groups. These abundant and evolutionarily ancient metabolites are found in nearly all living organisms^1^. The major polyamines in mammals are putrescine, spermidine, and spermine (Fig 1a), which are present at multimillimolar concentrations within cells^2^. Cells obtain polyamines through both *de novo* synthesis from the amino acids ornithine and methionine, as well as from dietary sources and cellular uptake^2^. These metabolites are essential for cell survival and polyamine depletion, either via genetic mutations or drugs, results in cytostasis and eventually cell death^3^.

**Fig. 1.**
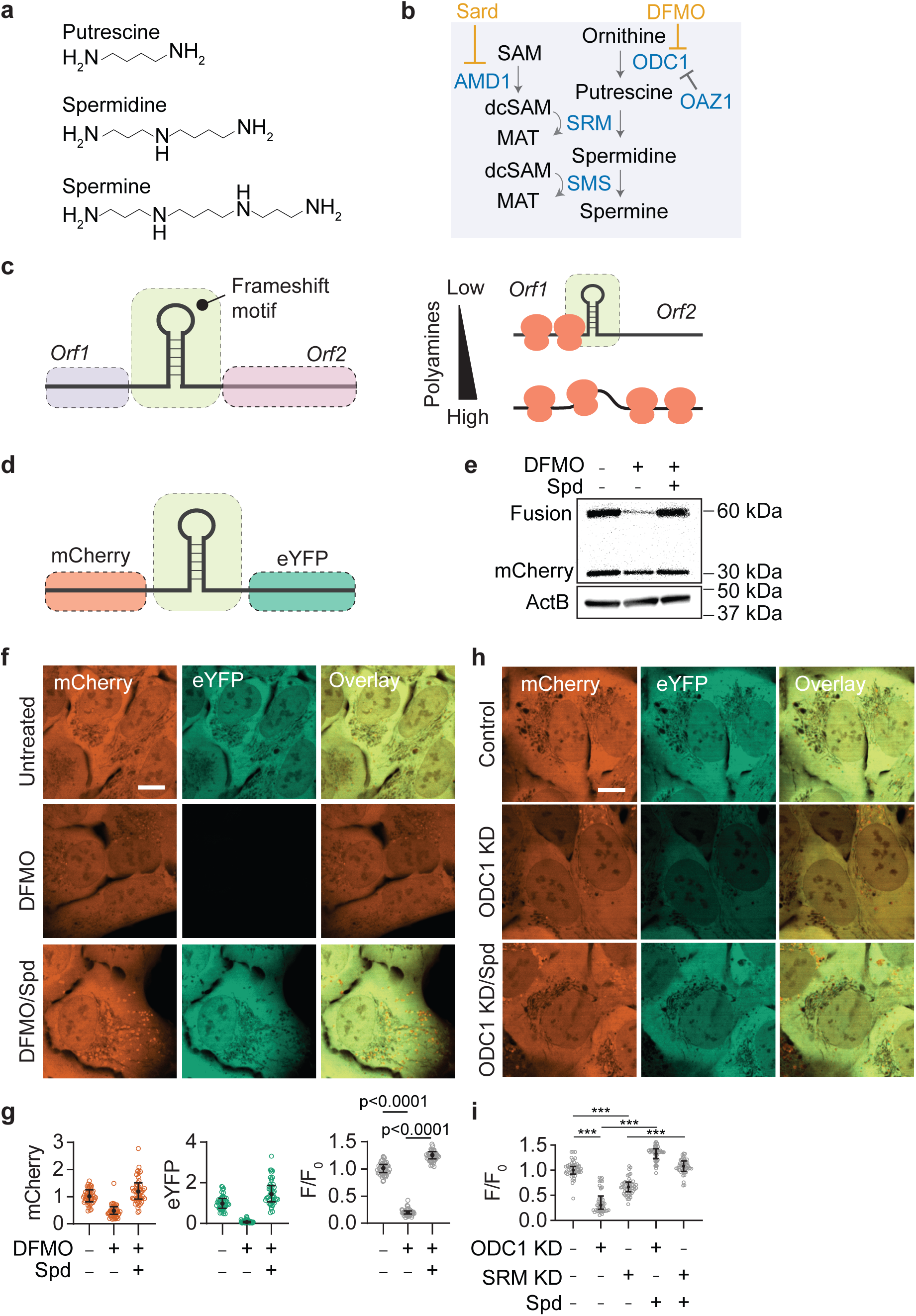
Polyamine sensor design and validation. **a,** Chemical structures of mammalian polyamines. **b,** Schematic of the polyamime biosynthesis pathway in mammalian cells. **c**, Schematic of polyamine-responsive +1 ribosomal frameshifting during translation of OAZ1 mRNA. **d-f,** Design for the polyamine sensor (d), corresponding immunoblot (e) and micrographs under indicated treatments. **g,** Quantification of frameshift efficiency using flow-cytometry for cells expressing polyamine sensor under indicated treatments. DFMO: difluoromethylornithine, 1 mM (90 h) and Spd: spermidine, 5 μM (18 h). **h-i,** Representative fluorescence micrographs (h) and corresponding quantification by flow cytometry of cells (i) upon genetic knockdown of indicated polyamine biosynthesis enzymes. Each data point shown in (g) and (i) represents a single cell. Error bars in (g) and (i) denote the median ± interquartile range and are calculated from ≥ 10000 cells. 50 data points are shown. Flow cytometry quantification and the fluorescent micrographs are representative of ≥2 independent experiments. Significance values are calculated using Student’s t-test. *** denotes p<0.0001. Scale bars, 10 µm.

At physiological pH, the amine groups of polyamines are protonated, enabling them to interact with and neutralize charge on various negatively charged molecules in the cell. Their distributed charge density allows polyamines to preferentially bind to nucleic acids, and a majority of the cellular polyamine pool is bound to RNA^4^. Perturbations to cellular polyamine levels affect numerous cellular processes, including chromatin organization, gene regulation, cell differentiation, autophagy, and immune function, underscoring the far-reaching impact of these small metabolites on cell physiology^2^. Polyamines bind to, and regulate ion channels, particularly inward rectifier potassium channels^5^ and NMDA receptors^6^, influencing membrane excitability and synaptic transmission. Spermidine also provides an amino-butyl group for an essential and evolutionarily conserved post-translational modification, hypusination, of the translation elongation factor, eIF5A^7^.

Given their myriad functions, both excess and deficiency of polyamines can lead to detrimental effects. Their cellular concentrations are tightly regulated through a complex network of feedback mechanisms that operate at the level of biosynthesis, catabolism, and transport^8^. A rate limiting enzyme in the biosynthesis of polyamines is ornithine decarboxylase (ODC1), which catalyzes the formation of putrescine (Fig. 1b). ODC1 levels are regulated by OAZ1, which binds to and promotes rapid ubiquitin-independent proteasomal degradation of ODC1, thus inhibiting polyamine synthesis (Fig. 1b)^9^. Increased polyamine levels also stimulate the production of catabolic enzymes such as SAT1, that acetylates polyamines for export^10^, and polyamine oxidases (PAOX and SMOX), which oxidize polyamines, generating hydrogen peroxide as a byproduct^11^. This production of reactive oxygen species underscores the cellular cost of polyamine dysregulation, as cells risk oxidative damage to prevent excess polyamine accumulation. Cells also modulate the polyamine transport pathways in response to intracellular polyamine concentrations, providing an additional level of feedback control^12^. The intricate interplay between these regulatory mechanisms ensures that polyamine levels are maintained within a narrow range optimal for cellular functions.

Consistent with the importance of maintaining cellular polyamine homeostasis, dysregulation of polyamine pathways is observed in cancer, aging, and neurological disease. Polyamine levels decline with age, and dietary polyamine supplementation in model organisms improves memory^13^, cardiac health^14^, reproductive aging^15^, and lifespan^16^. Mutations in polyamine regulatory enzymes are linked with an X-linked intellectual disability in Snyder-Robinson syndrome^17^ and with juvenile-onset Parkinson’s disease^18^. On the other hand, consistent with their roles in cell proliferation, the key polyamine biosynthetic enzymes as well as polyamine levels are elevated in many cancers driving cell transformation and tumour progression^19^.

Given the crucial role of polyamines in these pathological conditions, there is substantial interest in developing therapeutic strategies aimed at restoring polyamine balance^20,21^. However, although there is an overwhelming consensus that deviations in polyamine levels disrupt cellular functions, we currently lack technologies that allow us to measure polyamine concentrations at a cellular resolution. Current methods for detecting polyamines often rely on chromatography or mass spectrometry^18,22^. While such approaches have provided valuable insights into our understanding of polyamine homeostasis, they also have several inherent limitations. First, these methods offer only a snapshot of polyamine content averaged across the sample, losing information on cell-to-cell and temporal variability. Second, they require large sample quantities and have limited throughput, precluding large-scale genetic or pharmacological screens. Third, these methods require polyamine extraction from their native cellular context using organic solvents and extensive chemical derivatization^23^. These procedures introduce variability due to losses during extraction and labeling^24^, making it challenging to tease apart subtle differences.

Here we introduce a novel genetically encoded polyamine reporter that enables quantification of polyamine levels in living cells with single-cell resolution. Our reporter exploits the endogenous polyamine-responsive frameshift motif found in the OAZ1 gene, which plays a critical role in regulating cellular polyamine concentration. The OAZ1 mRNA contains a premature stop codon, and a programmed +1 ribosomal frameshift is required for the production of full-length, catalytically active OAZ1 protein (Fig. 1c). The efficiency of this frameshifting is dependent on the cellular polyamine levels, constituting an autoregulatory feedback mechanism to maintain polyamine homeostasis^25,26^. We leveraged the minimal polyamine-responsive frameshift module from OAZ1 and relayed this frameshifting efficiency to a fluorescent reporter, thereby generating a quantitative and live-cell readout for intracellular polyamine levels (Fig. 1d). This reporter allowed us to conduct a genome-wide screen to identify modulators of polyamine import and revealed an unexpected interaction between polyamine import and the mitochondrial respiratory complex.

## Results

### Sensor design and validation

We examined various segments of the OAZ1 gene and identified a 170-nucleotide region that retains polyamine-responsive frameshifting but lacks the catalytic domain (Supplementary Fig. 1a-h, Supplementary Note 1). This region was cloned between two fluorescent proteins, mCherry and eYFP (Fig. 1d). The eYFP sequence lacked an ATG start codon and was encoded in the +1-frame relative to mCherry. Ribosomes translating in the mCherry frame encounter a stop codon, and eYFP production requires a +1 frameshift. eYFP was chosen because its coding sequence lacks in-frame methionine codons within its N-terminus, and translation initiation at the encoded methionines should not produce a fluorescent protein^27^. The mCherry fluorescence signal allows normalization for cell-to-cell variations in transduction efficiency, mRNA stability, and cell’s translational status. The frameshift efficiency could be monitored via the relative fluorescence (F = eYFP/mCherry fluorescence) and reports on cellular polyamine levels. To prevent the accumulation of fluorescent proteins over time, which could confound the ratiometric readout, we expressed the reporter under a doxycycline-inducible promoter, ensuring that the fluorescence signal reports on the polyamine levels during a brief, well-defined time window.

Expression of this reporter in U-2OS cells resulted in both mCherry and eYFP proteins (Fig. 1e, f), demonstrating efficient ribosomal frameshifting. Western blots confirmed the production of the mCherry-eYFP fusion protein with a molecular weight consistent with a +1 ribosomal frameshift event (Fig. 1e). Insertion of a stop codon immediately after the mCherry coding region and before the OAZ1-derived frameshifting region effectively eliminated the eYFP fluorescence signal (Supplementary Fig. 2a-b). These results confirmed that eYFP expression is not due to alternative translation initiation within the eYFP sequence and that the eYFP signal does not arise from spectral bleed-through from the mCherry fluorescence. RNA sequencing analysis confirmed that the RNA transcribed from the reporter construct did not exhibit any unexpected transcriptional or post-transcriptional processing events that could account for eYFP production independent of frameshifting (Supplementary Fig. 2c). Reporter expression did not trigger measurable ODC1 degradation (Supplementary Fig. 2d) or alter cellular polyamine levels as measured by an ensemble assay (Supplementary Fig. 2e), ensuring that sensor expression does not appreciably perturb endogenous polyamine homeostasis.

Depletion of cellular polyamines using difluoromethylornithine (DFMO), a polyamine synthesis inhibitor^28^, reduced both eYFP and mCherry levels, consistent with the role of polyamines in protein translation^29^ (Fig. 1f, g). Notably, the decrease in eYFP signal was substantially more pronounced than that for mCherry (92% and 54% for eYFP and mCherry respectively, compared to the corresponding untreated controls). We quantified frameshifting efficiency as the ratio of eYFP to mCherry fluorescence (F), normalized to that for untreated cells (F_0_) to account for variations in fluorescence imaging conditions. This normalized ratio (F/F_0_) decreased five-fold (from 1.00 to 0.18) following DFMO treatment (1 mM for 90 h), but was restored (F/F_0_ ≈ 1.2) upon spermidine supplementation (5 μM spermidine for 18 h, Fig. 1g). Since polyamines can be oxidized by amine oxidases present in the serum, experiments involving polyamine supplementation and corresponding controls were performed in the presence of aminoguanidine, which potently inhibits amine oxidase activity^30^. Importantly, the stimulatory effect of polyamines on frameshifting was specific to the OAZ1 sequence. A control construct harboring the -1 ribosomal frameshift motif derived from SARS-CoV-2^31^ exhibited pronounced frameshifting under basal conditions, but its frameshifting efficiency was refractory to polyamine depletion (Supplementary Fig. 2f). Similar to DFMO treatment, genetic perturbations to the polyamine biosynthetic pathway, such as by ODC1 or SRM knockdown also reduced F/F_0_ for our reporter, which could be reversed by spermidine supplementation (Fig. 1h-i, Supplementary Fig. 2g). This polyamine dependent frameshifting (F/F_0_) was observed in other human cell lines (HEK293T, SH-SY5Y, and RPE1) as well as primary intestinal organoids derived from C57BL/6 mice (Supplementary Fig. 3a-e), demonstrating the versatility of our reporter and its potential for investigating polyamine biology across diverse cellular contexts. Taken together, these results demonstrate that our reporter system is sensitive to perturbations in cellular polyamine levels and can be used to monitor changes in endogenous polyamine homeostasis.

### Reporter calibration for polyamine quantification in live cells

Next, we calibrated the fluorescence readout against liquid chromatography-mass spectrometry (LC-MS) based polyamine quantification (see Reporter Calibration in Methods). We modulated the polyamine levels by titrating DFMO and compared the fluorescence readout from individual cells to ensemble polyamine measurements using LC-MS under equivalent treatments (Fig. 2a). This analysis showed that our reporter readout (F/F_0_), measured at single-cell resolution, correlated strongly with the average polyamine concentration measured by mass spectrometry (r^2^ = 0.79) (Supplementary Fig. 4a). Interestingly, F/F_0_ showed striking correspondence with the cellular spermidine concentration (r^2^ = 0.96) (Fig. 2b-d), but not with putrescine or spermine (Supplementary Fig. 4b). Similar results were obtained when cellular polyamine levels were changed by titrating sardomozide^32^, a polyamine biosynthesis inhibitor that targets AMD1 (Supplementary Fig. 4c-d). These observations are consistent with previous studies showing that the frameshifting in OAZ1 mRNA in cells is primarily stimulated by spermidine^33^. Although putrescine may also induce OAZ1 frameshifting, it requires ∼10-20-fold higher concentrations^34^ which are unlikely to be encountered under normal growth conditions (also see Supplementary Note 2). Likewise, while spermine can also promote OAZ1 frameshifting^34^, its biochemically available concentration is substantially lower (about 5-10 fold) than that of spermidine^4,35,36^. In agreement with this notion, prolonged treatment with a high dose of DFMO (500 μM for 3 days) reduced F/F_0_ to background levels (F/F_0_ = 0.02, representing a 50-fold decrease) despite unchanged spermine levels, as measured by mass spectrometry (Fig. 2c, Supplementary Fig. 4b). Collectively, these results emphasize that under normal physiological conditions, this sensor primarily reports on spermidine, which is the most abundant polyamine in mammalian cells^1^.

**Fig. 2.**
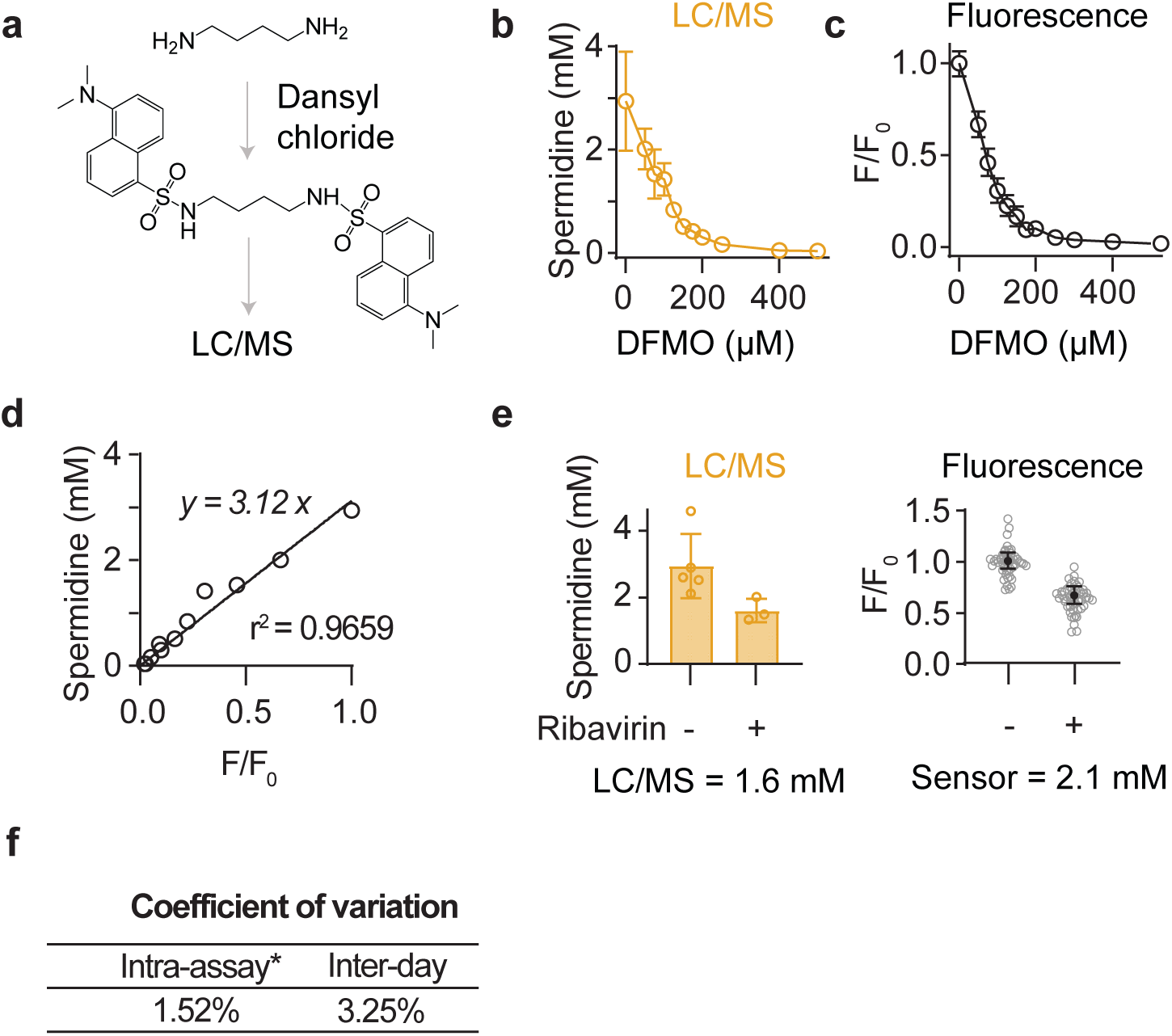
Reporter calibration and reproducibility. **a,** Schematic of LC/MS protocol for polyamine quantification. **b-c,** Intracellular spermidine concentration measured using LC/MS (b) and F/F_0_ (c) in response to various concentrations of DFMO. **d,** Linear regression of the cellular spermidine concentration with F/F_0_ (calculated from c). **e,** F/F_0_ and the spermidine concentration measured using LC/MS upon treatment with ribavirin (100 μM for 72 h; SAT1 activator). The sensor value was calculated using the fit from d. **f,** Coefficient of variation (CV) was calculated for intra-assay and inter-day variability. CV is calculated as the ratio of standard deviation to the average value. Error bars in (b) and (e, left) denote the mean ± standard deviation (n ≥3 independent experiments). Error bars in (c) and (e, right) denote the median ± interquartile range and are calculated from ≥ 10000 cells. 50 data points are shown.

The frameshift efficiency (F/F_0_) monotonically increased with cellular spermidine concentration, spanning a 100-fold range (from ∼40 μM to 4 mM) (Fig. 2d), underscoring the high sensitivity and large dynamic range of the reporter. To demonstrate that our reporter can effectively measure polyamine concentrations in live cells, we treated cells with ribavirin, a drug that was recently shown to deplete cellular polyamines by activating SAT1^22^. Remarkably, the cellular polyamine concentration estimated using our reporter closely matched those determined by conventional mass spectrometry-based analysis (Fig. 2e). Notably, while mass spectrometry required over ∼1 million cells, our reporter enabled measurements at single-cell resolution. Moreover, our reporter exhibited low technical variability, with a coefficient of variation of only ∼3% across multiple independent experiments (Fig. 2f, Supplementary Fig. 4e). In summation, these results show that the fluorescence readout from our reporter can be used to quantitatively measure cellular polyamine levels with high sensitivity and over a wide dynamic range in single living cells.

### Long-term tracking of polyamine dynamics in single cells

In the next steps, we sought to perform real-time monitoring of polyamine levels in individual cells over a longer duration. To further enhance the reporter’s performance for these longitudinal studies, we incorporated a degron tag^37^ to ensure rapid protein turnover, thus mitigating the reduction in dynamic range associated with the accumulation of fluorescent proteins (Fig. 3a). We also replaced eYFP with a near-infrared fluorescent protein, miRFP670-2^38^, to reduce the phototoxicity associated with long-term live-cell imaging. These modifications allowed us to track polyamine levels in cultured cells for >7 days with single-cell resolution (see Supplementary Note 3). Using this modified reporter, we examined the dynamics of polyamine depletion upon DFMO treatment and observed a gradual 50 ± 10% reduction in F/F_0_ over 3.5 days (Fig. 3b-d). Notably, there was considerable variability in how individual cells responded to DFMO, with some cells entirely evading its effects (Supplementary Fig. 4f), corroborating previous observations in clonal cells^39^. The reporter also enabled us to probe the kinetics of polyamine import. The addition of spermidine to DFMO-treated cells led to the restoration of polyamine levels over 2 days (Fig. 3b-d). Interestingly, we observed an initial overshoot of F/F_0_ relative to the baseline upon spermidine supplementation, which persisted for ∼40 h. This finding is consistent with recent observations that DFMO-treated cells are primed for polyamine import^40^ and underscores the role of feedback mechanisms in the regulation of polyamine homeostasis. Some cells showed an extended delay in transitioning out of the polyamine-low state upon spermidine addition (Supplementary Fig.4g). Collectively, these results demonstrate that our reporter provides a quantitative, near real-time measure of cellular polyamine concentration in single living cells. It reveals heterogeneity in drug response and provides a means to investigate molecular mechanisms of resistance and evasion.

**Fig. 3.**
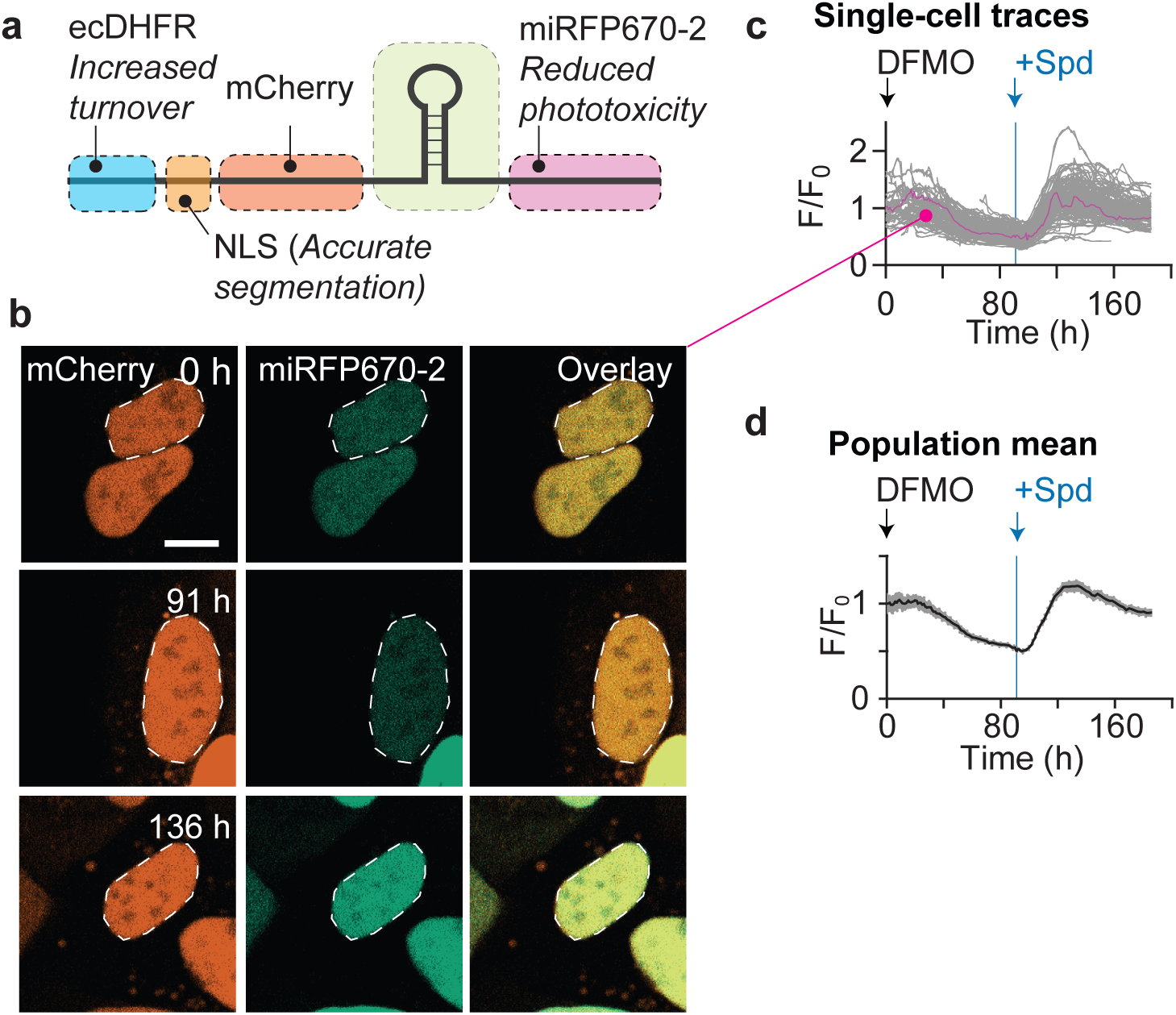
Live-cell longitudinal imaging of polyamine dynamics. **a,** Schematic of sensor design for real-time tracking of polyamines. **b,** Representative fluorescence images of cells expressing the degron-based sensor under indicated treatments. DFMO (2 mM), Spd (spermidine, 100 μM) and TMP (0.05 μM). Cells were grown in TMP starting 3 days before DFMO treatment to allow steady-state levels of the reporter expression. Media was replaced every two days with fresh reagents. **c,** Single-cell traces of F/F_0_ upon indicated treatments. F is the miRFP670-2 to mCherry fluorescence. F_0_ is the average frameshifting efficiency in untreated cells (at the time, t = 0). Arrows indicate the time-point of treatment. **d,** Mean F/F_0_ over time from the single-cell traces shown in c. Shaded bands depict the 95% confidence intervals.

### Genetic modifiers of polyamine import

Besides synthesis, mammalian cells can regulate their polyamine content by modulating transport but this pathway remains poorly understood^41^. One proposed polyamine import mechanism involves endosomal/lysosomal P5B-ATPases, ATP13A2 and/or ATP13A3^12^, that release lysosomal polyamines into the cytoplasm. Several other proteins have also been proposed to participate in polyamine uptake but their precise roles remain to be established^42–46^. Polyamine transport is also subject to elaborate feedback regulation, but the molecular factors involved in this process are not entirely elucidated^12^. We utilized our reporter to develop a polyamine import assay (Fig. 4a). In brief, cells were treated with DFMO to block polyamine biosynthesis, rendering them dependent on exogenously supplied spermidine. Polyamine import from the extracellular medium is expected to restore cellular polyamine concentration, which can be quantitatively read out using our reporter. As a proof-of-concept, we examined the effect of the drug AMXT-1501, a lipophilic polyamine mimetic that potently blocks polyamine transport^47^. Treatment with AMXT-1501 alone did not affect cellular polyamine levels (F/F_0_) indicating that under normal growth conditions, *de novo* polyamine synthesis is sufficient to maintain homeostatic levels (Fig. 4b-c). However, co-treatment of cells with AMXT-1501 and DFMO led to a striking 10-fold reduction in F/F_0_, which could not be rescued by exogenous spermidine supplementation (Fig. 4b-c). These results indicate that our assay allows us to monitor polyamine import in single cells, and can potentially help identify genetic and pharmacological modulators of this process.

**Fig. 4.**
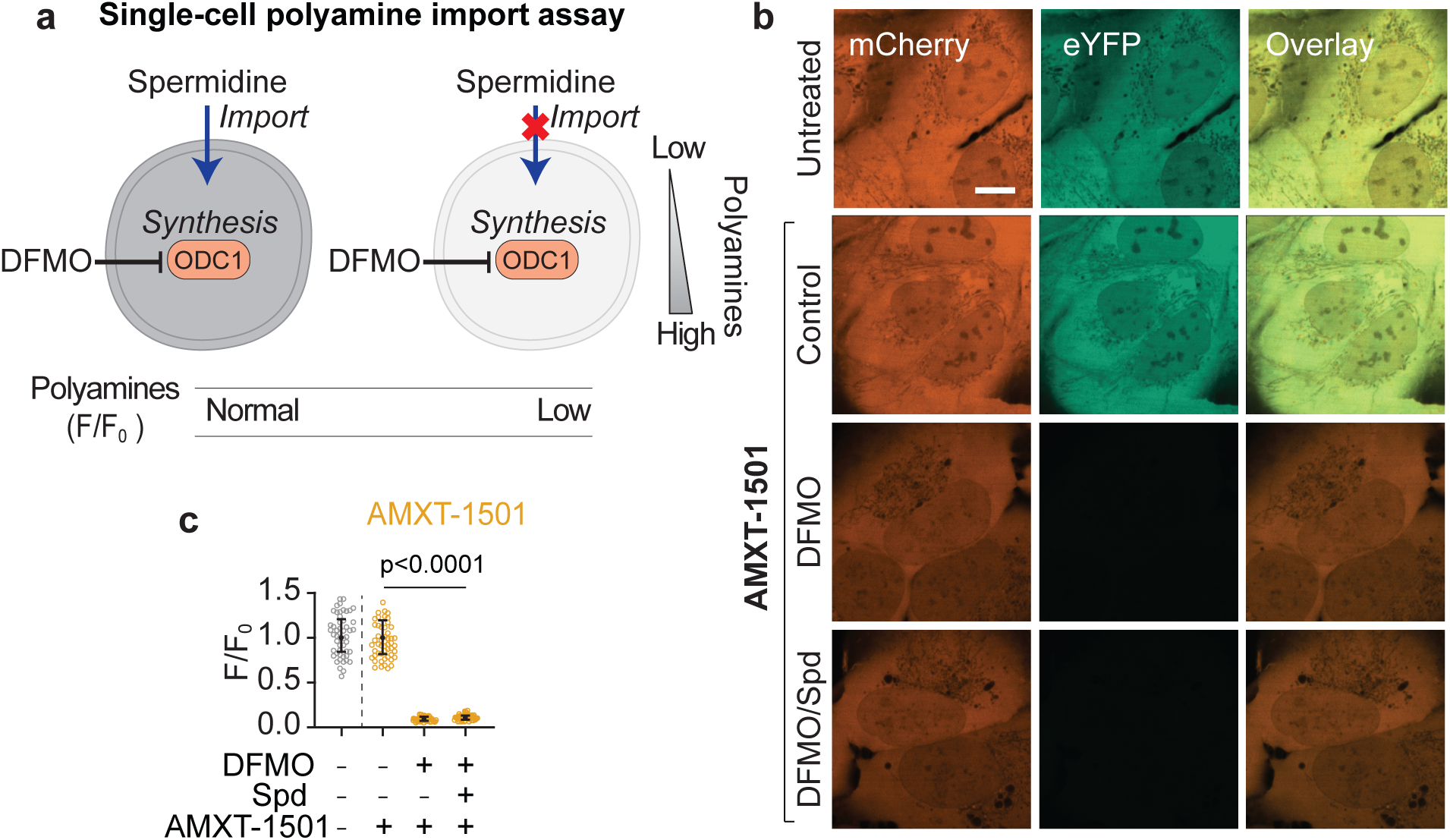
Single-cell polyamine import assay. **a,** Schematic of single-cell polyamine import assay. **b**, Representative fluorescence images of cells upon indicated treatments. AMXT-1501 (0.5 μM; polyamine import inhibitor), DFMO (1 mM), and spermidine (5 μM). **c,** Quantification of frameshift efficiency using flow-cytometry for treatments in (b).

Leveraging this assay, we performed a genome-wide CRISPR-Cas9 knockout screen to identify factors that influence polyamine uptake (Fig. 5a). We generated clonal K562 cells that express our polyamine sensor and transduced them with a pooled library of CRISPR single guide RNAs (sgRNAs) targeting 20,000 protein-coding genes with an average of 5 sgRNAs per target^48^. Following library transduction, cells were co-treated with DFMO and spermidine. Fluorescence-activated cell sorting (FACS) was used to isolate cells based on eYFP/mCherry fluorescence ratio. We expected that in the presence of DFMO and spermidine, import-competent cells would maintain their polyamine levels, while genetic knockouts that result in deficient spermidine import would have reduced polyamine content (Fig. 5a).

**Fig. 5.**
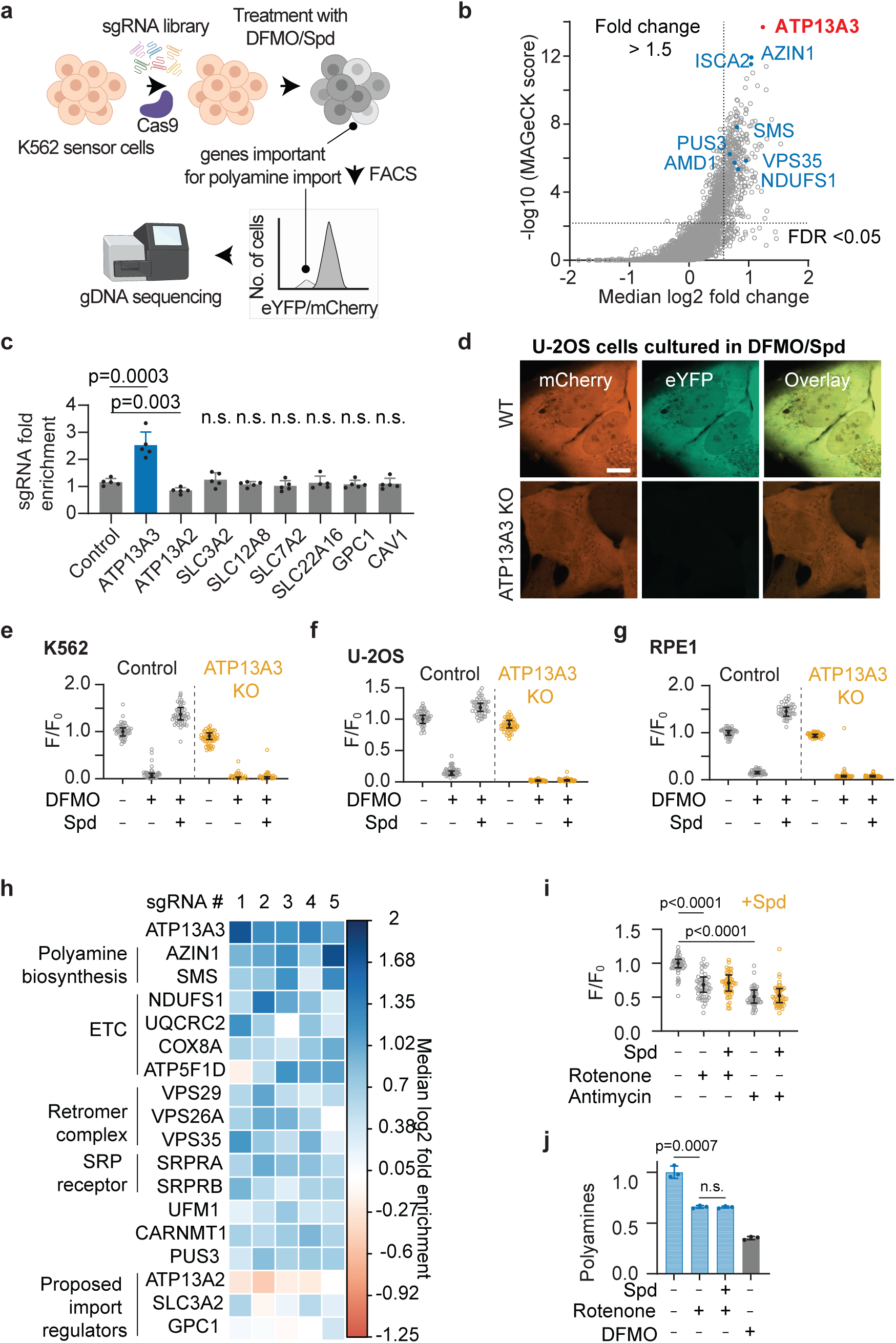
Genome-wide CRISPR screen identifies modifiers of polyamine import. **a,** Schematic of the CRISPR-Cas9 FACS-based genome-wide screen to identify modulators of polyamine import. **b**, -log10 (MaGeCK score) versus median log2 (fold enrichment) for all genes from the spermidine import screen. **c**, Graph depicting the fold enrichment for sgRNAs targeting indicated genes. Each data point is for a distinct sgRNA sequence. **d,** Representative images of U-2OS cells corresponding to treatment with DFMO: 1 mM and spermidine: 5 μM. (**e-g)** Comparison of polyamine levels in wild-type and upon ATP13A3 knock-out in K562 (e), U-2OS (f), and U-2OS (g) cells under indicated treatments. **h**, Heatmap of enrichment scores for individual sgRNAs targeting genes previously proposed to be involved in polyamine import. **i,** Comparison of polyamine levels in U-2OS cells under indicated treatments. Rotenone (1 μM; complex I inhibitor) and antimycin (1 μM; complex III inhibitor) were added 24 h before reporter induction and spermidine addition (5 μM, 18 h). **j**, Total polyamine levels (see enzymatic total polyamine assay in Methods) for the indicated samples (relative abundance). Rotenone (1 μM, 48 h) and spermidine (5 μM, 16 h). 2 mM DFMO (48 h) was used as a positive control. Error bars in (c) denote the mean ± standard deviation (5 independent sgRNAs). Error bars in (e-g, i) denote the median ± interquartile range and are calculated from ≥ 5000 cells. 50 data points are shown. Error bars in (j) denote the mean ± standard deviation from 3 independent experiments. Significance values are calculated using Student’s t-test. Flow cytometry quantification and the fluorescent micrographs are representative of ≥2 independent experiments. Scale bars, 10 µm.

ATP13A3, one of the known endosome-localized modulators of polyamine uptake, emerged as the top hit from the screen (Fig. 5b, Supplementary Table 1). Strikingly, knockout of ATP13A2, another P-type ATPase associated with polyamine import and Parkinson’s disease^49,50^, did not decrease polyamine import (Fig. 5c), despite being expressed at levels comparable to those of ATP13A3 in these cells (Supplementary Fig.5a). Moreover, knockout of other proposed polyamine transporters (for example, SLC3A2, SLC12A8, SLC7A2, and GPC1) had no discernible effect on polyamine import in our screen (Fig. 5c). Interestingly, ATP13A3 knockout was sufficient to completely abrogate spermidine uptake, phenocopying the effects of AMXT-1501 across multiple cell lines, including K562, U-2OS, and RPE1 cells (Fig. 5d-g, Supplementary Fig. 5b). The complete cessation of uptake upon ATP13A3 knockout demonstrates that it is an indispensable component of the spermidine import pathway in these cell types, and suggests that the release of polyamines stored in the endosomes may be the limiting step in controlling biochemically accessible polyamine levels in these cells.

Our screen also identified ∼180 putative modifiers of spermidine uptake (median log_2_ fold change > 0.75 with FDR < 0.05, Supplementary Table 1). These hits included known genes related to polyamine metabolism (Fig. 5h), including AZIN1 (antizyme inhibitor 1) and SMS (spermine synthase). The inhibition of polyamine import upon AZIN1 knockout is consistent with the known feedback repression of polyamine uptake caused by the stabilization of the antizyme family proteins upon AZIN1 depletion^51^. Likewise, SMS converts spermidine to spermine, and its loss leads to spermidine accumulation, which inhibits polyamine import^52^. Interestingly, several components of the mitochondrial electron transport chain (ETC) emerged as predominant modifiers of polyamine import (Supplementary Fig. 5c). To validate the effects of ETC inhibition on polyamine import, we treated U-2OS cells with pharmacological ETC inhibitors. Treatment with rotenone, a plant metabolite that potently inhibits complex I of the ETC, significantly decreased cellular polyamine levels (Fig. 5i). A similar decrease was observed following treatment with antimycin A, an inhibitor of complex III (Fig. 5i). Notably, reductions in polyamine levels due to rotenone or antimycin A treatment could not be compensated by exogenous spermidine supplementation, confirming import deficits (Fig. 5i, Supplementary Fig. 5d). This reduction in polyamine levels and deficient import was further verified using ensemble polyamine measurements (Fig. 5j). Other genes whose knockout reduced spermidine uptake included the subunits of the endoplasmic reticulum signal recognition particle (SRP) receptor (SRPRA and SRPRB), the retromer complex (VPS35, VPS26A, and VPS29), CARNMT1 (anserine synthase), PUS3 (a tRNA pseudouridine synthase), and UFM1 (involved in the UFMylation pathway) (Fig. 5h). Altogether, the ability to quantitatively monitor polyamines at single-cell resolution facilitated the identification of polyamine uptake modifiers and raises questions about the potential mechanisms linking mitochondrial respiration to polyamine transport.

## Discussion

We have established a novel genetically encoded fluorescent reporter that enables quantitative measurement of polyamine levels in living cells at single-cell resolution. This reporter system, based on the endogenous polyamine-responsive frameshift motif found in the OAZ1 gene, provides a real-time readout of intracellular polyamine concentrations with high sensitivity and a wide dynamic range. The simplicity of the assay greatly accelerates the measurement process, eliminating the need for extraction from native cellular context and extensive chemical derivatization. The single-cell resolution of the sensor enables high-throughput screens and may help identify new modulators of polyamine homeostasis. Moreover, the ability to track polyamine levels within the same sample could allow direct pharmacokinetic studies of polyamine-targeting therapies, including potentially in live animals.

Abnormalities in polyamine transport are implicated in several devastating diseases, including, Kufor-Rakeb syndrome (juvenile-onset parkinsonism with dementia)^49^, early-onset Parkinson’s disease^50^, amyotrophic lateral sclerosis (ALS)^53^, and pulmonary arterial hypertension^54^. Additionally, as rapidly proliferating cells exhibit high polyamine levels, there has been longstanding interest in developing polyamine inhibitors as cancer therapeutics^55,56^. Cancer cells can circumvent polyamine synthesis inhibition by increasing their uptake from the extracellular environment^19^, and there is a growing interest in targeting the polyamine import pathway via combination therapies^55,56^. Efforts to identify modulators of polyamine transport have been hampered by the lack of methods that provide a quantitative readout for polyamine uptake. Our sensor overcomes these technical limitations and may help elucidate the molecular details of polyamine transport and its regulation, potentially advancing therapy development.

Our genome-wide CRISPR knock-out screen established an essential role of ATP13A3 in polyamine import across different cell types and uncovered several new regulators of spermidine uptake. Strikingly, several of these polyamine import modifiers, such as VPS35 and subunits of ETC complex 1 (e.g., NDUFS1), are associated with Parkinson’s disease^57–59^. Likewise, rotenone exposure, which recapitulates key pathological and clinical features of the disease in animal models^60^, also affects polyamine uptake. These findings, along with the fact that loss of function mutations in the polyamine importer, ATP13A2, cause juvenile-onset Parkinson’s disease^18^, reinforce the link between polyamine homeostasis and the pathogenesis of this neurodegenerative disorder. Notably, recent work has shown that polyamine deprivation inhibits the synthesis of several mitochondrial proteins involved in oxidative phosphorylation^61^, suggesting that polyamine levels can influence mitochondrial function. Our work demonstrates the converse, that mitochondrial respiration, in turn, regulates polyamine homeostasis, highlighting a bidirectional cross-talk between these two processes.

One possible explanation for the inhibitory effects of ETC blockade on polyamine levels is the disruption of one-carbon metabolism. ETC inhibition impairs the tricarboxylic acid (TCA) cycle and leads to the depletion of aspartate^62,63^. This aspartate deficiency may activate ATF4^64^ and redirect one-carbon metabolism towards transsulfuration^65^, thereby limiting the availability of S-adenosylmethionine (SAM) for polyamine synthesis. ETC inhibition also suppresses *de novo* purine synthesis^66^. AMD1, a rate-limiting enzyme in polyamine synthesis, shares a strong co-dependency with both purine synthesis and oxidative phosphorylation enzymes (the Cancer Dependency Map^67^), suggesting a significant metabolic coupling between these pathways. We speculate that inhibition of polyamine biosynthesis in response to compromised ETC function is an adaptive response to suppress protein synthesis^29^ under energetically demanding conditions.

Polyamines are central to a multitude of cellular processes including gene expression, protein synthesis, cell proliferation, and cell death. There is growing interest in therapeutically harnessing this pathway but these efforts are hindered by gaps in our understanding of the molecular mechanisms involved in maintaining polyamine homeostasis. Polyamine regulatory pathways are also replete with intriguing biochemical regulations that defy common principles such as stop codon readthrough and frameshifting (in OAZ1)^25^, translation repression (of SAT1)^68^, ubiquitin-independent proteasomal degradation (of ODC1)^8^, and essential posttranslational modification, hypusination, unique to a single protein, eIF5A^7^. The availability of live-cell polyamine measurement reporters will provide a powerful platform to facilitate future discoveries into the biology of these abundant yet mysterious metabolites, advancing our understanding of disease mechanisms and potentially providing novel strategies for therapy development.

## Acknowledgments

We thank Miram Meziane, Kehui Xiang, Jimmy Ly, Mohamed El-Brolosy, Marine Krzisch, Norihiro Goto, Troy Whitfield, and the Jain lab members for helpful discussions. We are grateful to Chiara Alquati for assistance with organoid culture, Stephanie Long (USDA), Rakesh Minocha (USDA), and Subhash Minocha (UNH) for discussions on polyamine extraction, Keya Vishwanathan for help with CRISPR-Cas9 screen lentivirus preparation, and the Genome Technology Core at the Whitehead Institute for sequencing the libraries. We acknowledge the Whitehead Flow Cytometry Core Facility (Patrick Autissier, Aditya Rathee, and Kenji Goto-Hardy) for their assistance and Beverly Dobson for administrative support. This work is supported by grants from the NIH R35GM151111 (A.J.), Bumpus Foundation (A.J.), Pew Charitable Trusts (A.J.), and Chan Zuckerberg Initiative Collaborative Pairs Award (J.K.W. and A.J.).

## Author contributions

P.S. and A.J. conceived the project. P.S. designed, performed, analyzed, and interpreted all the experiments. C.Y.K. helped with the LC/MS instrument and analyzing related data with guidance from J.K.W.. H.R.K. helped with the CRISPR Cas9 screen and analyzing related data. S.I. generated and provided sensor-expressing organoids with guidance from O.Y.. A.B.J. helped with plasmid preparation. L.F. and T.K. established and optimized the protocol for LC/MS quantitation of polyamines in the lab. R.A. performed RNA sequencing analysis. A.J. supervised the project. P.S. wrote the manuscript with edits from A.J.. All authors reviewed the manuscript.

## Competing Interests

P.S. and A.J. are co-inventors on a pending patent application related to this polyamine reporter. J.K.W. is a member of the Scientific Advisory Board and a shareholder of DoubleRainbow Biosciences, Galixir, and Inari Agriculture, which develop biotechnologies related to natural products, drug discovery, and agriculture. All other authors have no competing interests.

## Online Methods

### Cloning and plasmid preparation

Complete plasmid sequences are provided in Supplementary Table 2. The sequence encoding the full-length human OAZ1 was obtained as double-stranded DNA fragments (IDT) and its various segments were cloned between PacI and BamHI sites in a doxycycline-inducible lentiviral transfer plasmid (pHR-Tre3G-mCherry-MCS-eYFP). pHR-Tre3G-mCherry-MCS-eYFP vector is derived by Gibson cloning from pHR-Tre3G-29xGGGGCC-12xMS2, a gift from Ron Vale (Addgene plasmid # 99149^69^). CRISPR-dCas9-mediated knockdown of ODC1 and SRM was performed using the plasmid pLV hU6-sgRNA hUbC-dCas9-KRAB-T2a-Puro (Addgene plasmid # 71236^70^), a gift from Charles Gersbach. Single-guide RNAs (sgRNAs) with high on-target efficiency to target ODC1 and SRM were designed as described previously^71^. The sgRNA sequences were as follows: ODC1 (GGGCGGCGGCGGCTACAGGA) and SRM (GGTAGCGCGAGCGCCGGTGG). CRISPR-Cas9 mediated knockout of ATP13A3 was performed using pLenti-EF1-SpCas9-FLAG-P2A-Puro-WPRE (a gift from Heather Keys) with sgRNA sequence TGGGTGAAATAACGAATCTG. For the live tracking experiments in Figure 1, the sequence encoding ecDHFR and miRFP670-2 were incorporated using Gibson cloning into a constitutive SFFV promoter-driven lentiviral transfer plasmid, pHR-tdMS2CP-YFP-WPRE (Addgene 99151^69^), a gift from Ron Vale. A plasmid containing ecDHFR sequence was a gift from Iain Cheeseman and miRFP670-2 was amplified from pTubulin-miRFP670-2 (Addgene 197238^38^), a gift from Kiryl Piatkevich. All cloning and plasmid preparations were performed in Stbl3 *Escherichia coli* cells (Invitrogen, C7373-03) grown at 30°C. All plasmids were sequence verified by Sanger or Oxford Nanopore sequencing (Plasmidsaurus or Quintara Bioscience).

### Chemicals

The following reagents were purchased from Sigma-Aldrich: perchloric acid (244252), 1,7-diamino heptane (D17408), acetone (270725), toluene (34866), L-proline (P0380), acetonitrile (34851), sodium carbonate, dansyl chloride (311155), spermidine trihydrochloride (S2501), DMSO (276855), and aminoguanidine hydrochloride (396494). DL-α-difluoromethylornithine (DFMO; 16889), rotenone (13995), antimycin A complex (34799), and ribavirin (16757) were obtained from Cayman Chemicals. Sardomozide dihydrochloride (HY-13746B) and AMXT-1501 (HY-124617A) were ordered from MedChemExpress.

#### Preparation of compounds and inhibitors

Fresh stocks were prepared for all the amine compounds from powder before the experiments. Polyamines and aminoguanidine stocks were prepared to a final concentration of 100 mg/mL in PBS for each experiment. Sardomozide was dissolved in DMSO to final concentrations of 10 mM. DFMO (55 mM) and ribavirin (100 mM) stocks were prepared in water. All stocks were filtered through a 0.45 µm filter upon preparation and stored at -20℃.

### Cell culture

U-2OS, HEK 293T, K562, RPE1 (immortalized), and SH-SY5Y were purchased from ATCC and authenticated by short tandem repeat profiling. U-2OS, HEK 293T, and RPE1 were propagated in DMEM (Gibco 11965126). Suspension cultures of K562 cells were cultured in RPMI-1640 (Gibco 11875093). SH-SY5Y cells were grown in EMEM (ATCC 30-2003)/F12 (Gibco 11765054). All media were supplemented with penicillin/streptomycin/glutamine (Gibco 10378016) and 10% (v/v) tetracycline-free fetal bovine serum (FBS) (Gibco, 26140079), except for SH-SY5Y cells which had additional FBS (15% v/v). Cells were maintained at 37°C with 5% CO_2_ in a humidified incubator. Cells were tested for mycoplasma contamination.

### Lentivirus preparation and transduction

We first generated cell lines stably expressing reverse tetracycline transactivator (rtTA) protein (Clontech) via lentiviral transduction. These stable cell lines were then transduced with the polyamine reporter under a tetracycline-inducible promoter. Lentivirus was generated as follows. The lentivirus transfer plasmids (2 µg), the packaging plasmid psPAX2 (1 µg), and the envelope plasmid pCMV-VSV-G (0.5 µg) were mixed with 8 µL of lipofectamine LTX (Invitrogen, 15338-100) in 500 µL of Opti-MEM reduced serum media (Gibco, 31985-070) for transfection of HEK293T cells according to the manufacturer’s recommended protocol. After two days, the lentiviral particles were collected by passing the supernatant through a 0.45-μm filter and used to transduce cells supplemented with 10 μg/mL polybrene (Millipore Sigma TR1003G). Polybrene was excluded during the transduction of SH-SY5Y cells due to toxicity. Transgene expression of the polyamine sensor was induced by adding 1000 ng/mL doxycycline (Sigma-Aldrich) for 18 h except where otherwise specified. The various indicated treatments (DFMO, rotenone, etc.) were retained during this induction period.

### Crypt isolation and intestinal organoid culture

As previously reported^72^ and briefly summarized here, small intestines were removed, washed with cold PBS, opened longitudinally, and then incubated on a shaker at 4℃ with PBS plus EDTA (10 mM) for 45 min. Tissues were then moved to PBS. Crypts were mechanically separated from the connective tissue by shaking and then filtered through a 70-μm mesh into a 50 mL conical tube to remove villus material and tissue fragments. Isolated crypts for cultures were embedded in Matrigel (Corning 356231, growth factor reduced) and cultured in a modified form of the medium as described previously^73,74^. Crypt culture media consists of Advanced DMEM (Gibco 12491015) that was supplemented with EGF 40 ng ml^-1^ (PeproTech 315-09), Noggin 200 ng ml^-1^ (PeproTech 250-38), R-spondin 500 ng ml^-1^ (Peprotech 315-32), B27 1X (Life Technologies 17504044), CHIR99021 3 µM (LC laboratories C-6556), and Y-27632 dihydrochloride monohydrate 10 µM (Sigma-Aldrich Y0503). Matrigel was allowed to solidify for 20-30 minutes in a 37℃ incubator. The culture medium was then overlaid onto the Matrigel, changed every three days, and maintained at 37℃ in fully humidified chambers containing 5% CO_2_.

To generate the polyamine sensor-organoids, organoids derived from mouse intestines were dissociated by Triton-X at 37℃ for 2 min, mixed with virus supernatant (prepared as described above) and polybrene at 9 µg/ml, and then transferred to 48 well plates. The plate was centrifuged at 600× g at 30℃ for 1 h, and incubated in chambers for 3 h. Collected samples were embedded in Matrigel and incubated with crypt medium plus 5% FBS for 7 days. 1000 ng/mL of doxycycline was added into the medium and organoids were incubated for 24 hours to express the fluorescent protein, and then fluorescent-positive organoids were picked up under the fluorescent scope. For DFMO and spermidine treatment, organoids were treated after passaging. 1 mM DFMO with or without 50 µM spermidine was added to the culture medium, and were imaged 3 days after treatment. The commonly used organoid medium supplement B27 contains polyamines, and thus differences in the polyamine levels upon DFMO treatment become apparent only in its absence.

### Enzymatic polyamine assay

For Fig. 1f and Fig. 6f, measurements were performed using the fluorescent quantification assays (abcam239728) as per the manufacturer’s instructions. The polyamine assay buffer (no added polyamine standard) was used as the background.

### Microscopy and image analysis

Cells were seeded on a glass-bottom 96-well plate (Brooks, MGB096-1-2-LG-L) and imaged on a Dragonfly 505 spinning-disk confocal microscope (Andor Technologies) with a piezo Z-stage (ASI), an iXon Ultra 888 EMCCD camera and appropriate filter sets. Organoids were cultured in polymer-coated chamber slides (ibidi 80286). Pin-hole size was kept at 40 μm. Z-stacks were acquired with a step size of 0.3 to 1 µm depending on the thickness of the sample using a 100× oil immersion objective NA 1.45 (Nikon, MRD01905, pixel size 121 nm × 121 nm). During the imaging, cells were kept in a humidified chamber (OKO labs) maintained at 37° C and 5% (v/v) CO_2_. The following laser excitation and bandpass emission filters were used: YFP (488 nm, 521/38 nm), mCherry (561 nm, 594/43 nm), and miRFP670-2 (637 nm, 620/60 nm). For each experimental condition, at least 40 randomly chosen cells were imaged in at least two independent replicates. Representative images were generated by taking an average projection of three slices centered at the focused plane.

For the live-cell polyamine tracking in Fig. 3, we quantified the fluorescence signal in the microscopy images at each time point. F_0_ is miRFP670-2/mCherry at t=0, before DFMO treatment A custom FIJI script was used to find the focal plane of Z-stacks using a normalized variance algorithm (https://imagejdocu.list.lu/macro/autofocus_hyperstack). Background fluorescence was computed from the average signal of 10 regions devoid of any cells. This background was calculated for each channel and subtracted before intensity quantification. Cells were segmented from background-corrected mCherry fluorescence images using Cellpose^75^, a deep learning-based segmentation algorithm (a pre-trained “cyto” model was directly applied without additional training). Segmented objects with a diameter less than 25 pixels were excluded from downstream analysis. TrackMate^76^ was used to perform tracking of segmented cells employing a Linear Assignment Problem (LAP) framework which uses a Jonker-Volgenant solver. In brief, a cost matrix was generated for frame-to-frame linking using a maximal allowed frame-to-frame linking distance of 75 pixels and with linking cost as the distance squared. Next, the built track segments are linked using the LAP framework again, allowing the branching of a track into two sub-tracks to account for mitosis events with a maximal distance of 75 pixels. Gap-closing was also included to create a link over two spots separated by a missed Cellpose detection with a maximal distance of 150 pixels over 2 frames. These parameters were optimized by manually benchmarking the predicted cell tracks on a subset of the data. Only the longest track was retained for each identified trajectory. Cells entering or leaving the field of view were retained for analysis. Lineages with frames less than half of the total number of frames during the entire experiment duration were excluded. Incorrect segmentation or tracking events were manually filtered out. Custom MATLAB scripts were used to process the single-cell integrated intensity data. For the images shown in Fig. 3, we cropped regions containing a particular tracked single-cell and used the background corrected image. The raw intensity values for miRFP60-2/mCherry were normalized using an exponential fit derived from a control, untreated sample imaged under similar conditions to account for the loss in fluorescence signal due to photobleaching.

### Flow cytometry

For fluorescence quantification, the cells were analyzed by flow cytometry (BD LSR II, BD FACSDiva). At least 5,000 cells were analyzed for each experimental condition in at least two independent replicates unless otherwise noted. The flow cytometry data was then analyzed in FlowJo (TreeStar). Background fluorescence was determined for each channel in each experiment from the average signal of uninduced samples, and was subtracted before intensity quantification. GraphPad Prism was used to perform statistics and generate plots. Cells underwent the same treatment as for microscopy.

### Western blotting

Cells were washed with ice-cold PBS and lysed with RIPA lysis buffer (25 mM Tris-HCl pH 7.5 (Invitrogen 15567027), 150 mM NaCl (Invitrogen AM9760G), 1% (v/v) NP-40 (Fisher Scientific AAJ19628AP), 1% (w/v) sodium deoxycholate (Sigma-Aldrich D6750), 0.1% (w/v) SDS (Bio-Rad 1610302)) supplemented with 1% (v/v) HALT protease and phosphatase inhibitors (Thermo Scientific 78429) and 125 U/mL Benzonase nuclease (EMD Millipore E1014). The lysate was homogenized and incubated on ice for 30 min with vortexing every 10 min. Cell debris was removed by centrifugation at 21000× g for 10 min at 4°C and lysates were mixed with 4× Bolt lithium dodecyl sulfate (LDS) sample buffer (Invitrogen B0007) with 100 mM DTT (Thermo Scientific R0861) and heated at 70°C for 5 min. Samples were separated on a Bolt 4-12% Bis-tris polyacrylamide gel (Invitrogen, NW04122) and then transferred to a PVDF membrane (Invitrogen IB24002) using an iBlot 2 dry blotting system (Invitrogen IB21001). Membranes were blocked in 5% (w/v) nonfat dry skim milk (BD Biosciences 232100) in TBST (tris-buffered saline (Fisher Scientific AAJ60764K3) with 0.1% (v/v) Tween-20 (Fisher Scientific BP337) for 1 h at room temperature. Membranes were incubated with primary antibodies diluted in 1% (w/v) skim milk in TBST at 4°C overnight. After 4X TBST washes, membranes were incubated with appropriate secondary antibodies in 1% (w/v) skim milk in TBST for 1 h at room temperature. Membranes were washed 4X with TBST, and chemiluminescence signals were detected using SuperSignal West Femto Maximum Sensitivity Substrate (Thermo Scientific 34095) with a ChemiDoc XRS+ imager (Bio-Rad). The antibody details are as follows: rabbit anti-mCherry (Rockland, 600-406-379, 1:5000), mouse HRP anti-beta-actin (ab20272, 1:50,000), anti-ODC1 (ab193338, 1:1000) and goat anti-rabbit HRP conjugate (Sigma A0545, 1:5000).

### RNA sequencing and splicing analysis

Total RNA was extracted with PureLink RNA Mini Kit (Invitrogen) per the manufacturer’s instruction. PolyA selection and RNA library construction were performed by Novogene with sequencing as 150×150 base paired-end libraries using an Illumina NovaSeq 6000 (Novogene), to a depth of ≥ 20M reads. Reads were aligned to the human genome (hg38, annotation file GRCh GTF 38.93) and plasmid map for the reporter plasmid using the short-read alignment tool STAR (version 2.7.1a)^77^ with the default options. Sashimi plots for splicing analysis of the reporter were generated with ggsashimi (version 1.1.5)^78^.

### Polyamine extraction for LC/MS

This protocol was adapted from previous work^23^. In brief, two million K562 cells were washed twice with ice-cold PBS, pelleted, and then liquid nitrogen was used to quench metabolism. The samples were transferred temporarily to dry ice before being stored in a sealed container at -80°C. The tubes were then thawed in 400 μL 0.2 M perchloric acid (PCA) and resuspended. 10 μL of 0.1 mg/mL 1,7 diamino heptane (in PCA, prepared fresh) was spiked in as a non-biological polyamine internal standard. After a brief incubation on ice (15 min) and intermediate vortexing, the cell lysate was cleared by centrifugation at 21,000 xg for 10 min at 4°C, and 100 μL of the supernatant was neutralized with 200 μL of 3 M sodium carbonate (in water). Finally, 400 μL of 7 mg/mL dansyl chloride (in acetone) was added. The solution was mixed by vortexing and incubated at 60°C for 30 minutes (protected from light from now on). To quench the excess dansyl chloride, 100 μL of 100 mg/mL L-proline (in water) was added to the tube and mixed by vortexing. After 10 minutes of incubation at room temperature in the dark, the top phase (acetone solution) was completely removed and discarded. Next, 500 μL of toluene was added to the tube, and the solution was mixed by vortexing. We extracted dansylated polyamines into toluene because, in the aqueous phase, they stick to the walls of the polypropylene microtubes and do not efficiently get transferred. After a brief incubation at room temperature (15 minutes), the tube was cleared by centrifugation at 11,000 xg for 1 min, and 200 μL of the toluene phase (top) was aliquoted into a 9 mm wide autosampler glass vial, and then dried using a Mulitvap nitrogen evaporator under dark conditions. Needles were continuously adjusted to expedite the drying process. The samples were stored at -80°C until the day of analysis. Standard solutions of polyamines (all three mixed together) were prepared at the concentrations of 0.0002, 0.0004, 0.001, 0.002, 0.005, 0.01, 0.02, 0.05, 0.1, 0.2, 0.5, 1, 2, 5, 10, 20, 50, 100, 200, 500 and 1000 μM and were treated with the same procedure as the sample. All reagents were prepared fresh immediately before use, and all the experiments were performed in at least three replicates. Due to the large number of samples, we performed metabolite extractions in multiple batches (16 individual samples at a time). To mitigate analytical bias in downstream analysis, the replicates of an experimental condition were distributed in different batches.

### Polyamine LC/MS analysis and quantification

On the analysis day, the dried samples were centrifuged to collect all samples to the bottom of the tube, resuspended in 20 μL acetonitrile, vortexed for 5 minutes, and then spun down again before placing them into the autosampler. All the steps were performed at 4°C, and exposure to light was avoided. Liquid chromatography was conducted on a Vanquish Flex Binary UHPLC system (Thermo Fisher Scientific) using water with 0.1% formic acid as solvent A and acetonitrile with 0.1% formic acid as solvent B. Reverse phase separation of analytes was performed on a ZORBAX RRHT Eclipse XDB-C18 column, 4.6 x 50 mm, 80 Å pore size, 1.8 µm particle size (Agilent, No. 927975-902; guard column, Agilent, No. 820750-903). The column oven was held at 25°C, and the injection volume was 2 µL. All injections were eluted with 70% B for 2.0 min, a gradient of 70-90% B for 1.0 min, 90-95% B for 1.0 min, 95-97.5% B for 2.0 min, 97.5-70% B for 1.0 min and 70% B for 5.0 min, with a flow rate of 1 mL/min. All mass spectrometry analyses were performed on a high-resolution Orbitrap Exploris 120 benchtop mass spectrometer (Thermo Fischer Scientific) operated in positive ionization mode with a full scan range of 550-1150 *m/z* and top four data-dependent MS/MS scans. The orbitrap resolution is 120,000 with RF lens of 70% and static spray voltage of 3500 V. Single-ion monitoring scans were also collected along with each method for targeted detection of the following compounds– 597.2565 *m/z* [M+H]^+^ at 3.3 min; dansyl heptane, 555.2126 *m/z* [M+H]^+^ at 2.0 min; dansyl putrescine, 845.3163 *m/z* [M+H]^+^ at 4.0 min; dansyl spermidine, and 1135.427 *m/z* [M+H]^+^ at 5.0 min; dansyl spermine– with a scan width of 5.0 *m/z* and RT window of 1.0 min. All MS data were collected under the profile data type. Multiple acetonitrile blank and polyamine standards were interspersed throughout the run to track any technical drifts in MS signal quality.

Chromeleon 7.2.10 ES, TSQ Tune 3.1.279.9, and XCalibur 4.5 (Thermo Fisher Scientific) were used for deconvolution, peak alignment, identification, and quantitation of the raw file peaks. The .raw files were imported onto MZmine 3 for quantitative analysis of dansyl heptane, dansyl putrescine, dansyl spermidine, and dansyl spermine. The following MZmine 3 modules were used: Mass detection, FTMS shoulder peak filter, ADAP chromatogram builder, Local minimum feature resolver, 13C isotope filter, Join aligner, and Same RT and *m/z* range gap filter. The MZmine 3 batch analysis file containing all processing parameters is attached as a .xml file in Source Data. The peak area for each polyamine was normalized by the peak area of the internal standard to represent the feature (polyamine) abundance before visualization and statistical analysis on GraphPad Prism (v9.5.1).

### Reporter calibration

In brief, we determined the polyamine concentrations in cells by quantifying the moles of polyamines using LC/MS-based analysis and the cell volume by confocal microscopy. To estimate the cell volume, cells were mounted and imaged in the mCherry channel. Z-stacks with a step size of 0.25 μm were acquired and the total cell volume was estimated using a custom CellProfiler pipeline, as has been previously reported (https://github.com/CellProfiler/tutorials/tree/master/3d_monolayer). At least 50 cells were analyzed and the cell culture conditions were kept identical to those used for LC/MS analysis. The mean cell volume within each condition was then used. For LC/MS analysis, a standard curve was prepared using individual polyamines. The metabolite abundances in extracts were obtained by comparing the LC/MS data against standards fit to a linear equation. The total number of moles of a polyamine in the cell extract was then calculated from the sample concentration and the corresponding sample volume. Cellular polyamine concentrations were calculated using the total moles of a metabolite in a sample, the total number of cells per sample, and the average volume of each cell.

The exact value of eYFP/mCherry is dependent on the laser settings of the flow cytometer, and thus we opted to calibrate the polyamine concentrations using the fold change (F/F_0_) rather than the absolute value (F). To estimate the absolute cellular spermidine concentrations (mM) from fluorescence (F/F_0_), we interpolated the data from DFMO titration to create a regression curve where the x-value is the F/F_0_ and the y-value is the absolute spermidine concentration (mM). Fluorescence was then used to determine the approximate spermidine for ribavirin treatment. Polyamines are commonly added to commercial kits for RNA transcription and translation and hence, it was not feasible to perform these calibrations using in vitro translation of the sensor mRNA.

### Lentivirus production for CRISPR-Cas9 screen

15 x 10^6^ HEK-293T cells were seeded in T175 cm^2^ flasks in DMEM (Thermo Fisher Scientific #12430054) supplemented with 10% fetal bovine serum (GeminiBio #100-106). After 24 hours, the media was changed to 20 mL viral production medium: IMDM (Thermo Fisher Scientific #1244053) supplemented with 20% inactivated fetal serum (GeminiBio #100-106). At 32 hours post-seeding, cells were transfected with a mix containing 76.8 µL Xtremegene-9 transfection reagent (Sigma Aldrich #06365779001), 3.62 µg pCMV-VSV-G (Addgene plasmid # 8454)^79^, 8.28 µg psPAX2 (a gift from Didier Trono; Addgene plasmid # 12260), and 20 µg sgRNA/Cas9 plasmid and Opti-MEM (Thermo Fisher Scientific #11058021) to a final volume of 1 mL. Media was changed 16 hours later to 55 mL fresh viral production medium. The virus was collected at 48 hours post-transfection and filtered through a 0.45 µm filter, aliquoted, and stored at -80 °C until use.

### CRISPR-Cas9 screen

A genome-wide lentiviral sgRNA library^48^ in a Cas9-containing vector comprising 97,888 unique sgRNA sequences with ∼5 sgRNAs per target (Supplementary Table 3) was used to transduce 600 x 10^6^ K562 cells to achieve an MOI < 1 (30-50% transduction efficiency) and ∼1000-fold library coverage.

Briefly, polybrene (10 µg/mL final concentration) and virus were mixed with cells (2.5 x 10^6^ cells/mL final density) and distributed into individual wells in 6-well plates. Plates were centrifuged at 1126 x *g* for 45 minutes at 37 °C and transferred to an incubator. After 8 hours, cells were pelleted, the virus was removed, cells were resuspended in the fresh growth medium, and transferred to T225 cm^2^ flasks (250,000 cells/mL final density). After 36 hours, cells were collected and reseeded in fresh growth medium (200,000 cells/mL final density) and puromycin was added (3 µg/mL final concentration). After 3 days, cells were collected and transduction efficiency was determined by comparison of cell survival of transduced cells relative to control cells (untransduced and unselected). Cells were passaged every 2 days (0.2 x 10^6^ cells/mL) for 3 days before reporter induction using doxycycline for 24 hours. At the screen endpoint, cell pellets were collected from flasks representing 1000-fold library coverage of unsorted cells and frozen at –80 °C. Induced cells were sorted and cell pellets were collected.

### Sequencing library preparation

Genomic DNA (gDNA) was extracted from cell pellets of 40-50 x 10^6^ cells using the QIAamp DNA Blood Maxiprep Kit (Qiagen # 51192) according to manufacturer’s instructions with minor modifications: QIAGEN Protease was replaced with 500 µL of a 10 mg/mL solution of ProteinaseK (MilliporeSigma # 3115879001) in water; cells were lysed overnight; centrifugation steps after Buffer AW1 and AW2 were performed for 2 minutes and 5 minutes, respectively; 1 mL of water preheated to 70 °C was used to elute gDNA (5-minute incubation), followed by centrifugation for 5 minutes. gDNA was quantified using the Qubit dsDNA HS Assay kit (Thermo Fisher Scientific #Q32851).

All PCR reactions were performed in 50 µL reactions using ExTaq Polymerase (Takara Bio #RR001B) using the following primers:

Forward: 5’- AATGATACGGCGACCACCGAGATCTACACCCCACTGACGGGCACCGGA - 3’

Reverse: 5’- CAAGCAGAAGACGGCATACGAGATCnnnnnnTTTCTTGGGTAGTTTGCAGTTTT - 3’

Where “nnnnnn” denotes the barcode used for multiplexing.

For all samples, 1, 3, or 6 µg of gDNA was initially amplified for 28 cycles in 50 µL test PCR reactions. Subsequently, an additional 23.33 reactions were performed using 6 µg per reaction (140 µg gDNA). Reactions were pooled and 100 µL of each reaction was purified using Select-a-Size DNA Clean and Concentrator (Zymo Research #D4080), eluted with 15 µL water, and quantified using the Qubit dsDNA HS Assay kit before sequencing for 50 cycles on an Illumina Hiseq 2500 using the following primers:

Read 1 sequencing primer: 5’- GTTGATAACGGACTAGCCTTATTTAAACTTGCTATGCTGTTTCCAGCATAGCTCTTAAAC - 3’

Index sequencing primer: 5’- TTTCAAGTTACGGTAAGCATATGATAGTCCATTTTAAAACATAATTTTAAAACTGCAAACTACCCAAGAAA - 3’

### CRISPR screen data analysis

Sequencing reads were trimmed and mapped to the sgRNA library using Bowtie version 1.0.0^80^ and counted. Data was analyzed using MAGeCK version 0.5.9.5^81^ with the following parameters: gene test false discovery rate (FDR) threshold of 0.05; FDR method for p-value adjustment; median as the gene-level scoring metric; sgRNAs targeting intergenic regions as control sgRNAs. Pre-ranked gene set enrichment analysis (GSEA)^82^ was performed with version 4.1.0 using the Human_Gene_Symbol_with_Remapping_MSigDB.v2023.1.Hs chip and the c2.cp.kegg_medicus.v2023.2.Hs.symbols (c2.cp.wikipathways.v2023.2.Hs.symbols^83^) gene set. The enrichment statistic was set to classic, normalization was set to meandiv, and the maximum score was used as the collapsing mode. Data was visualized using R version 4.2.1 corrplot package version 0.92^84^ and base graphics, and GraphPad Prism version 10.

### Statistics and reproducibility

Statistical details can be found in the figure legends for each figure.

## Reporting Summary

### Data availability

Source data is available from the corresponding author upon request.

## Notes

### Summary of Updates

Introduction and Figures 2-4 were revised for clarity.

