## Supplementary text for "Genetically encoded fluorescent reporter for polyamines"

**Supplementary Information**

**Supplementary Note 1**

**Design and characterization of the polyamine reporter**

The human *OAZ1* gene consists of 685 coding nucleotides with an in-frame UGA stop codon located at the nucleotides 205-207 (NCBI NM_004152.3). Production of full-length catalytically active OAZ1 requires a +1 ribosomal frameshift at the stop codon. The sequence spanning 1-275 nucleotides in OAZ1 mRNA was used as the starting point for reporter design as it does not bind ODC1, lacks the catalytically active C-terminal domain^16,17,67^, but is expected to retain the frameshift activity. This truncated version contains a 60-nucleotide long frameshift stimulatory RNA pseudoknot three nucleotides downstream of the slippery site (UCC UGA), potentially acting as a mechanical stressor to force the translating ribosome into a new reading frame^68^. The second mutant contained only nucleotides 106-275, cloned downstream of an ATG start codon. Previous ribosome profiling experiments have shown that a sizable fraction of ribosomes are paused in this region, suggesting a putative role in translational repression^69^. This construct (106-275) also avoids potential interactions of the translated N-terminal OAZ1 protein with the mitochondrial import machinery^70^. A third construct (185-275) consisting of the last few codons in ORF1, the frameshift site, and pseudoknot was cloned based on the evolutionary conservation of the nucleotide sequence 5’ to the shift site across homologs of vertebrates antizyme genes^71^. We also generated a mutant lacking the pseudoknot (1-216) to measure its contribution to polyamine-mediated frameshifting. These regions were individually cloned between mCherry and YFP fluorescent proteins, under a doxycycline-inducible promoter. Each construct was stably integrated into U-2OS cells and was evaluated for 1) its ability to induce ribosomal frameshift, and 2) its responsiveness to cellular polyamine levels as determined by reduction in frameshifting upon treatment with DFMO (an irreversible inhibitor of ODC1) and its rescue after adding exogenous spermidine (Supplementary Fig. 1). All constructs retained the ability to induce polyamine-responsive frameshift, albeit to varying degrees. The presence of the pseudoknot significantly enhanced the frameshifting efficiency. The 106-275 mutant was chosen for the polyamine sensor since this sequence is relatively short, retains high frameshifting efficiency, and produces a substantial change in the eYFP:mCherry fluorescence ratio between DFMO treatment and spermidine supplementation.

**Supplementary Note 2**

Titrations with sardomozide, an inhibitor of AMD1 that leads to excessive putrescine accumulation while depleting spermidine and spermine, revealed that F/F_0_ correlated with intracellular spermidine content but not with putrescine (Supplementary Fig. 4c-d). When cells were treated with a relatively high dose of sardomozide (500 nM), the cellular putrescine levels substantially increased by ~25-fold, but the frameshift efficiency, F/F_0_, decreased to 0.35 (F/F_0_ = 1 for untreated cells). These results suggest that putrescine does not efficiently stimulate +1 ribosomal frameshifting on OAZ1 mRNA, which is consistent with a previous study indicating that putrescine can only induce comparable +1 ribosomal frameshifting on OAZ1 at concentrations ~10-20 fold higher than spermidine^25^. This observation also explains the higher offset in F/F_0_ observed in sardomozide-treated cells compared to what would be expected based on DFMO titrations. Therefore, under typical physiological conditions, putrescine is likely to only have a modest effect on our reporter readout.

**Supplementary Note 3**

**Measuring temporal changes in intracellular polyamine levels**

A key advantage of our reporter is that it allows one to assess changes in polyamine levels within the same cell over time. We engineered a set of additional features in the reporter to enable long-term live-cell imaging (Fig. 3a): (1) First, we replaced eYFP with the far-red miRFP670-2 to reduce phototoxicity associated with long-term blue light illumination. miRFP670-2 also lacks in-frame methionine codons within its N-terminus and has minimal spectral overlap with mCherry. (2) We used the DHFR (dihydrofolate reductase) tag for protein destabilization^29^ to improve the response kinetics. The small molecule ligand, trimethoprim (TMP), was constitutively added to stabilize the DHFR fusion protein. TMP concentration was empirically optimized to facilitate rapid turnover of the fusion protein while maintaining an appropriate signal-to-noise ratio. (3) We incorporated a nuclear localization signal in the reporter to aid cell segmentation for automated tracking by demarcating the nucleus. These modifications allowed us to monitor polyamine levels in live cells in response to the drug DFMO over 8 days using time-lapse microscopy. We observed a continuous decline in the frameshift efficiency, F/F_0_ (F = miRFP670-2/mCherry, F_0_ is the mean F at t = 0), after DFMO treatment which lasted for ~90 h until it reached a steady low value of F/F_0 ­_≈ 0.5 (Fig. 3c-d). Spermidine addition to cells in this low-polyamine state triggered an increase in F/F_0_, observable within ~6 h after supplementation. At around 40 h after spermidine addition, F/F_0_ peaked at a value ≈ 1.2, about 20% higher than that observed in untreated cells. Subsequently, F/F_0_ gradually started returning to the basal levels and stabilized after ~30 h. These data indicate that inhibition of polyamine biosynthesis using DFMO elicits a compensatory increase in polyamine uptake, and results in a temporary excess. The increased uptake is under negative feedback regulation, leading to a gradual return to basal homeostatic levels. Our single-cell analysis also revealed significant cell-to-cell heterogeneity in this response. A small subset of cells in the population escaped the effects of DFMO for as long as ~90 h (Supplementary Fig. 4f). Some cells showed an extended delay in transitioning out of the polyamine-low state upon spermidine addition (Supplementary Fig. 4g). Note that the F/F_0_ in these experiments (measured using miRFP670-2 and mCherry fluorescence) is not directly comparable to the results in Fig. 1/2 due to differences in fluorescent proteins and protein turnover kinetics.

**Legends for Supplementary Data Figures**

**Supplementary Fig. 1 | a-d,** Representative fluorescent micrographs of cells expressing various OAZ1-derived candidate sequences under indicated treatments. DFMO (1 mM; 90 h) and spermidine (5 μM; 18 h). The position of the nucleotide sequence in the endogenous OAZ1 mRNA transcript is labeled in blue, where the start codon is encoded by bases 1-3. **e-h,** Quantification by flow cytometry for samples in b-e. Each data point represents a single cell. Error bars denote the median ± interquartile range and are calculated from ≥ 5000 cells. 50 data points are shown. Flow cytometry quantification and the fluorescent micrographs are representative of ≥2 independent experiments. Significance values are calculated using Student’s t-test. Scale bars, 10 µm.

**Supplementary Fig. 2 | a-b,** Representative fluorescence images (a) and corresponding flow cytometry quantification (b) of cells expressing either the polyamine sensor or its mutated version with a stop codon immediately after the mCherry coding region. **c,** RNA sequencing analysis and Sashimi plot for cells expressing the sensor. The x-axis is the base coordinate within the sensor construct, and the y-axis indicates the number of reads per million mapped sequencing reads. Splicing events with at least 10 reads per junction, as detected by the STAR aligner^66^, are indicated by arcs. Spliced sensor transcripts represent a negligible population (approximately 0.06%) of the total, and would not affect the F/F_0_. WPRE: Woodchuck hepatitis virus Posttranscriptional Regulatory Element. Each data point shown in (b) represents a single cell. Error bars denote the median ± interquartile range and are calculated from ≥ 5000 cells. 50 data points are shown. Flow cytometry quantification and the fluorescent micrographs are representative of ≥2 independent experiments. Significance values in (b) are calculated using Student’s t-test. Scale bars, 10 µm. **d,** Immunoblot of the indicated samples using an anti-ODC1 antibody. **e,** Total polyamine levels (see Enzymatic total polyamine assay in Methods) for the indicated samples (in nmol per million cells). DFMO (2 mM for 48 h) was used as a positive control. **f,** Representative fluorescence images and corresponding flow cytometry quantification of cells expressing a reporter construct containing a -1 ribosomal frameshift motif derived from the SARS CoV2 genome under indicated treatments. DFMO (1 mM; 90 h) and spermidine (5 μM; 18 h). **g,** Representative fluorescence images of cells expressing our polyamine reporter with SRM knock-down and after supplementation with spermidine (5 μM; 18 h). Each data point shown in (b) and (f) represents a single cell. Error bars in (b) and (f) denote the median ± interquartile range and are calculated from ≥ 5000 cells. 50 data points are shown. Flow cytometry quantification and the fluorescent micrographs are representative of ≥2 independent experiments. Significance values are calculated using Student’s t-test. Scale bars, 10 µm.

**Supplementary Fig. 3 | a-d,** Representative fluorescence images and corresponding quantification of frameshift efficiency in HEK293T (a), SH-SY5Y (b), RPE1 (c) cells, and primary murine intestinal organoids (d) expressing sensor under indicated treatments. We observed two distinct populations of cells in intestinal organoids, out of which, only one population responded to DFMO treatment and subsequent spermidine supplementation. The other population exhibited a low F/F_0_ value suggesting low polyamine biosynthesis and import. Each data point represents a single cell. Error bars denote the median ± interquartile range and are calculated from ≥ 5000 cells. 50 data points (cells) are shown for representation purposes. Flow cytometry quantification and the fluorescent micrographs are representative of ≥2 independent experiments. **e,** Flow-cytometry quantification of eYFP and mCherry for organoids in (d). 500 data points are shown. Significance values in (a)-(c) are calculated using Student’s t-test. Significance values in (d) are calculated using the Mann-Whitney non-parametric test. Scale bars denote 10 µm in (a)-(c) and 50 µm in (d).

**Supplementary Fig. 4 | a,** Linear regression of the total polyamine concentration with F/F_0_ (calculated from DFMO titration). **b,** Intracellular putrescine (purple) and spermine (blue) concentration measured using LC/MS in response to DFMO titration. **c,** Intracellular spermidine concentration measured using LC/MS (yellow) and F/F_0_ (grey) in response to sardomozide titration. **d,** Intracellular putrescine (purple) and spermine (blue) concentration measured using LC/MS in response to sardomozide titration. **e,** Quantification of frameshift efficiency using flow-cytometry in cells expressing polyamine sensor in three independent experiments over three days (nine independent experiments). F_0_ is the ratio of eYFP to mCherry fluorescence in cells on Day 1 (replicate 1). All replicates are plotted on the graph. Error bars in (c, grey) and (e) denote the median ± interquartile range and are calculated from ≥ 5000 cells. 50 data points are shown. Error bars in (b), (c, yellow) and (d) denote the mean ± standard deviation (n ≥3 independent experiments). **f-g,** Cell-to-cell heterogeneity in response to DFMO treatment and subsequent spermidine rescue: sample cell whose polyamine levels are unresponsive to DFMO treatment (f) and a cell where the maximal spermidine uptake is delayed by 60 h (g).

**Supplementary Fig. 5 | a,** DESeq2 normalized read counts from RNA-sequencing of K562 cells. **b,** Representative fluorescence micrographs of cells in Fig. 6c. **c,** Barplot of -log10(FDR) for gene sets (WikiPathways GSEA enrichment) with positive FDR < 0.05 (requiring > 2 sgRNAs/gene) with abbreviated gene set names. **d,** Comparison of polyamine levels in U-2OS cells under indicated treatments. DFMO (1 mM), rotenone (1 μM; complex I inhibitor), and antimycin (1 μM; complex III inhibitor) were added 24 h before reporter induction and spermidine addition (5 μM, 18 h. Error bars in (a) denote the mean ± standard deviation from 3 independent experiments. Error bars in (d) denote the median ± interquartile range and are calculated from ≥ 5000 cells. 50 data points are shown. Flow cytometry quantification and the fluorescent micrographs represent ≥2 independent experiments. Significance values are calculated using Student’s t-test. Scale bars, 10 µm.

**Legends for Supplementary Tables.**

**Supplementary Table 1.** Guide RNA enrichment scores from genome-wide CRISPR-Cas9 screen for spermidine import.

**Supplementary Table 2.** Complete sequences of the plasmids used in this study.

**Supplementary Table 3.** Sequences of the Cas9 sgRNAs and their corresponding targets used for the spermidine import screen.
