## Supplementary figures for "Genetically encoded fluorescent reporter for polyamines"

Suppl. Fig. 1| Selection of a minimal sequence for the polyamine sensor

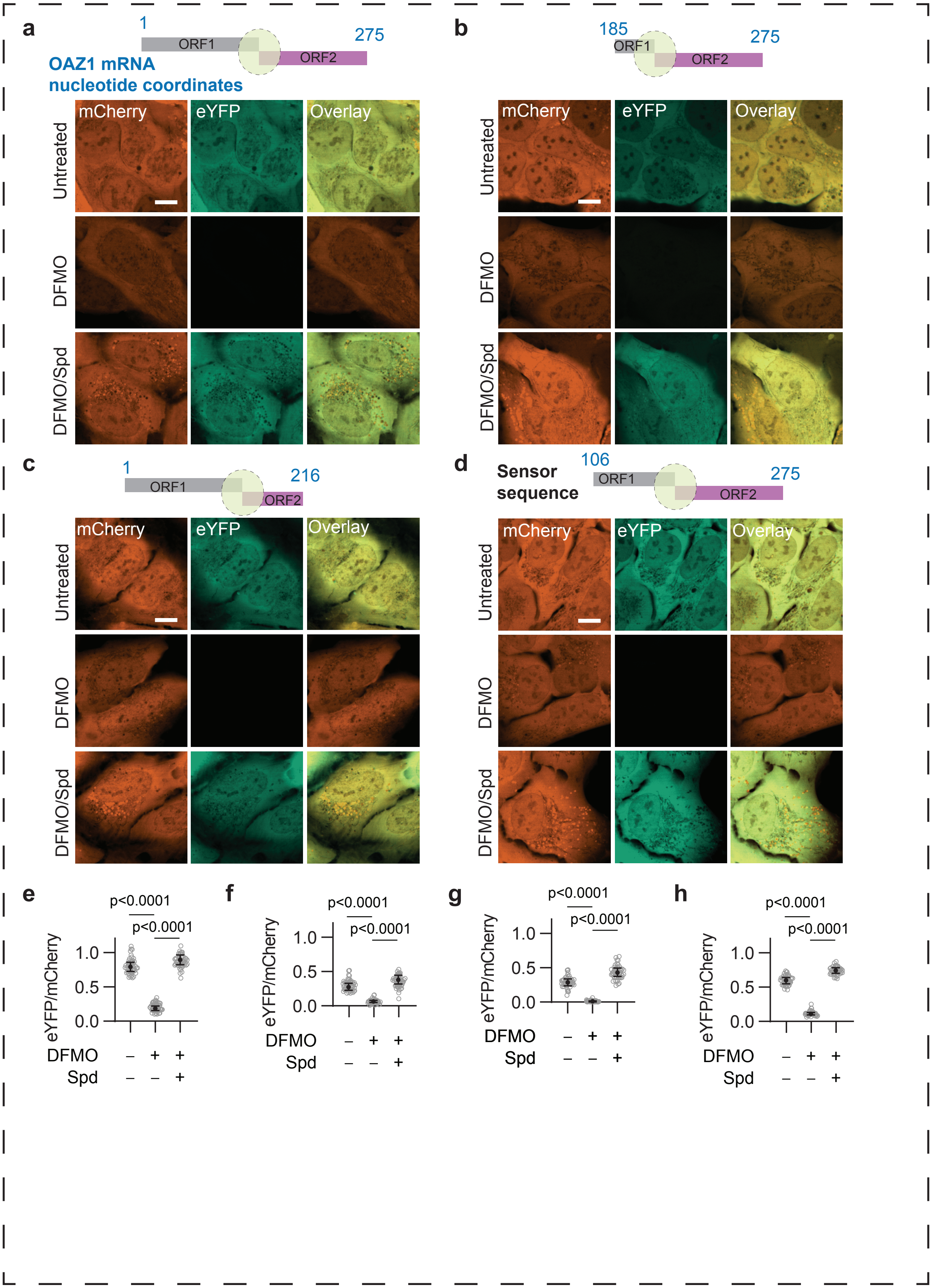

### Suppl. Fig. 2 | eYFP expression is driven by a +1 ribosomal frameshifting event and is independent of cryptic post-transcriptional events

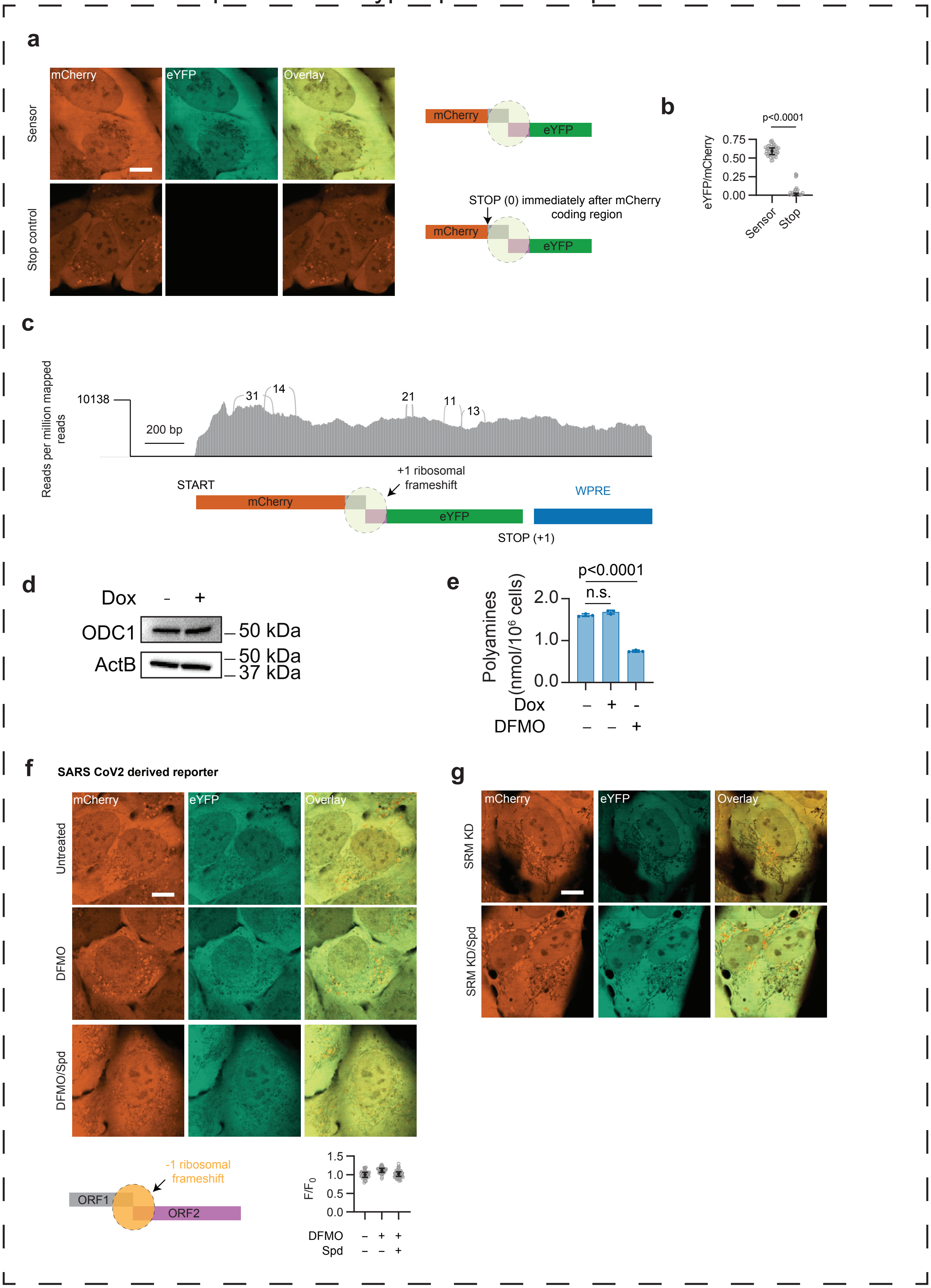

### Suppl. Fig. 3 | Sensor can measure polyamines in multiple cell types and organoids

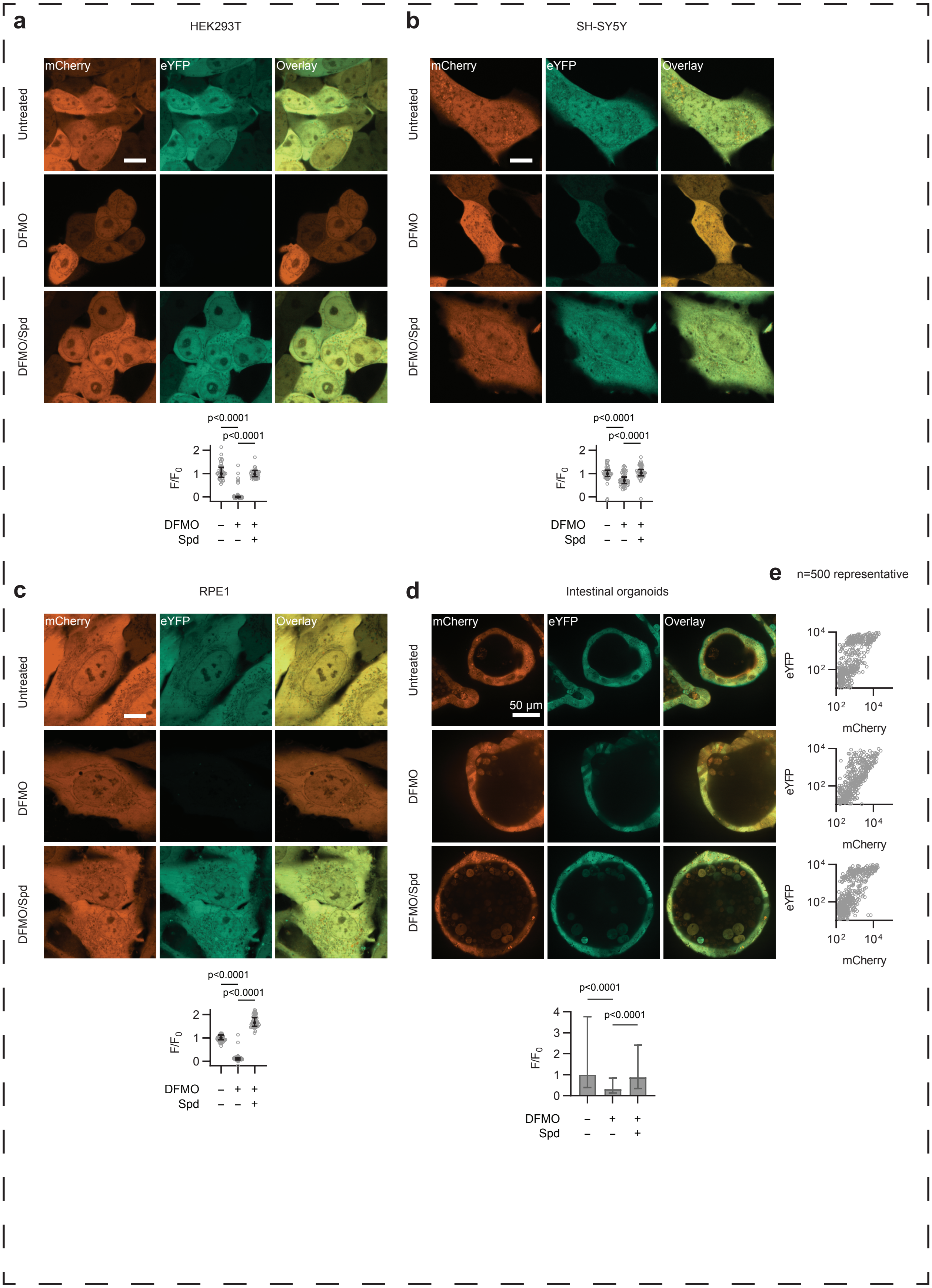

### Suppl. Fig. 4. Sensor primarily reports on spermidine levels and allows longitudinal tracking in single cells

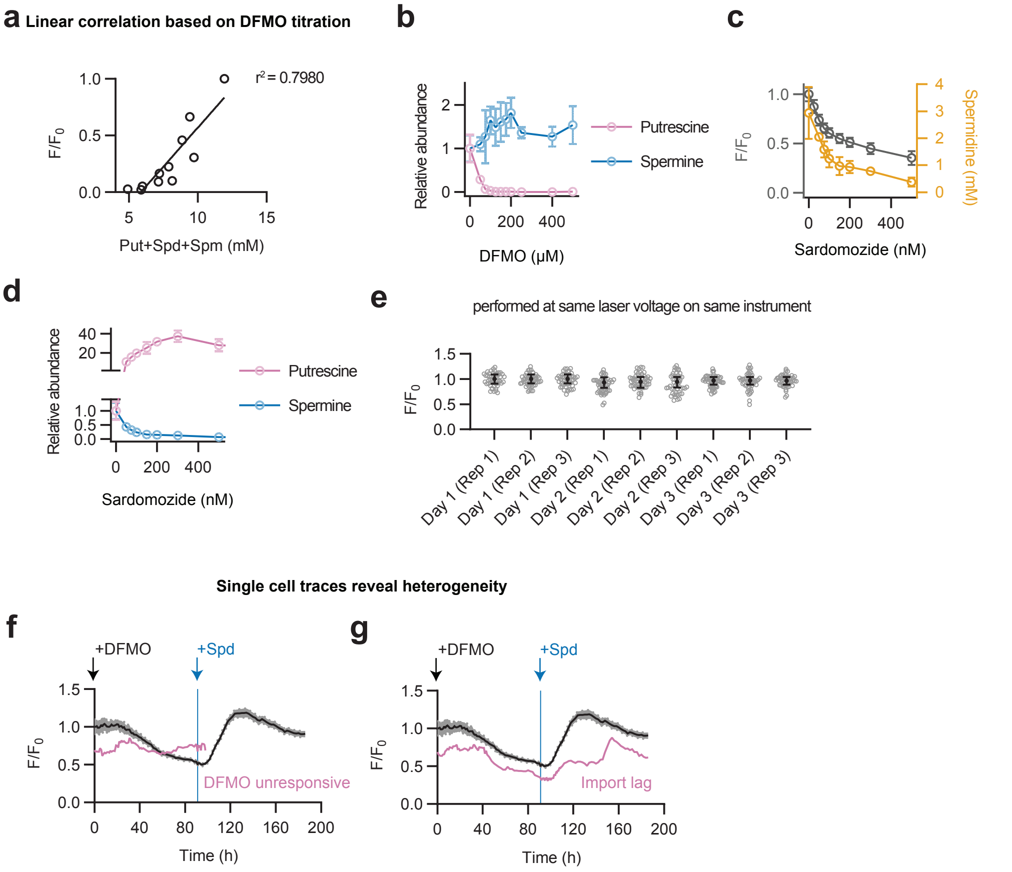

Suppl. Fig. 5. Polyamine import regulation and gene set enrichment analysis under metabolic inhibition

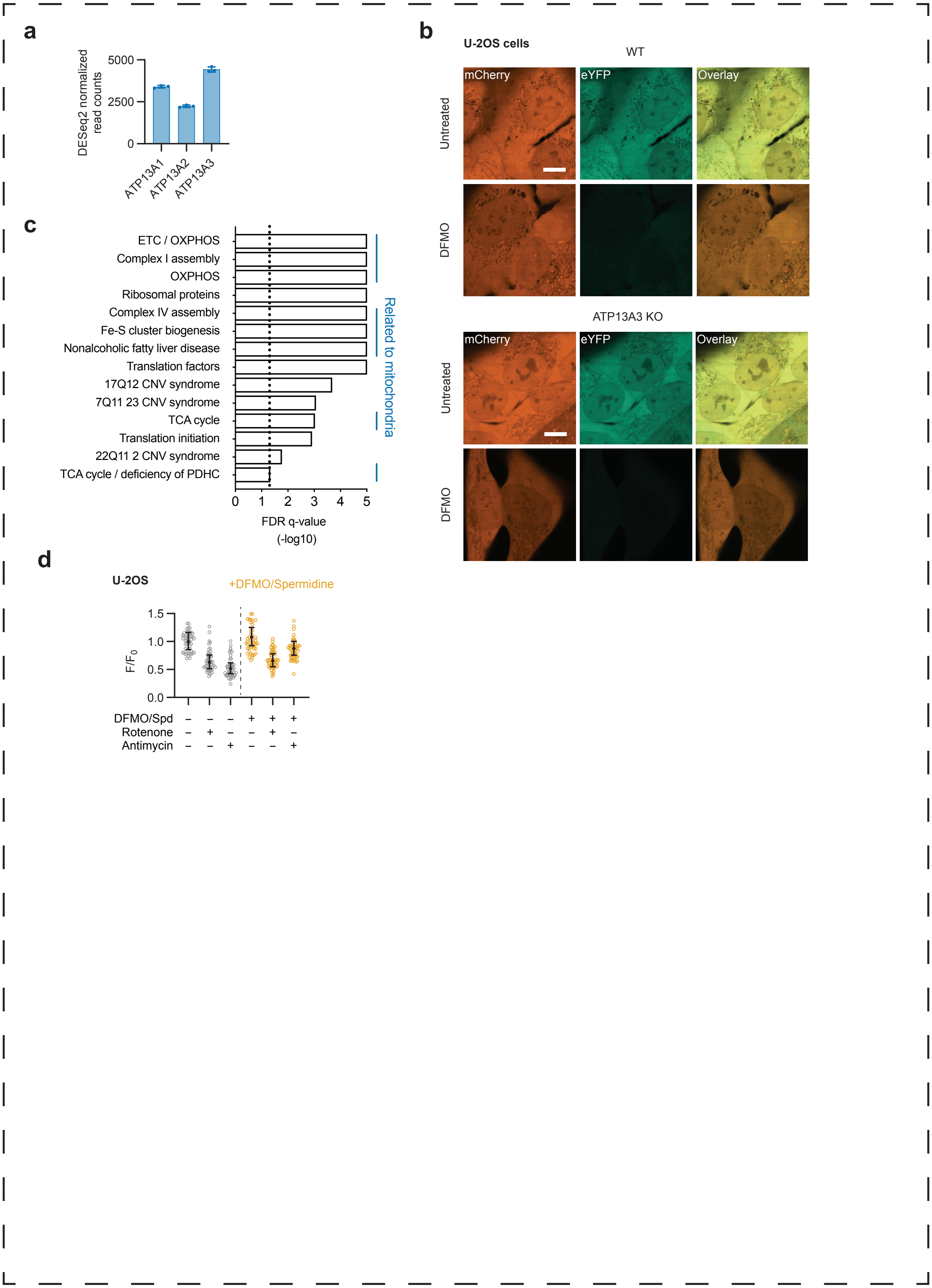
